# EDSpliCE, a CRISPR-Cas9 gene editing platform to rescue splicing, effectively corrects inherited retinal dystrophy-associated splicing defects

**DOI:** 10.1101/2024.03.27.587013

**Authors:** Pietro De Angeli, Stefanida Shliaga, Arturo Flores-Tufiño, Eleonora Roschi, Salome Spaag, Katarina Stingl, Laura Kühlewein, Bernd Wissinger, Susanne Kohl

## Abstract

**Background:** Correct splicing of transcripts is essential to ensure the production of functional gene products in eukaryotic cells. Missplicing of transcripts has been identified as the underlying molecular mechanisms behind various disease-causing variants in a wide range of inherited genetic conditions. Achieving therapeutic splicing correction is possible through antisense oligonucleotide and CRISPR/Cas9 strategies. However, while antisense oligonucleotides offer effective modulation, they do not enable for permanent correction. On the other hand, current CRISPR/Cas9 approaches often rely on dual-gRNA-inducing deletion of larger pieces of DNA, containing the site(s) responsible for the splicing defect, particularly the elimination of pseudoexons, raising concerns about potential chromosomal instability.

**Results:** The novel gene editing strategy, Enhanced-Deletion Splicing Correction Editing (EDSpliCE), just uses single gRNAs to effectively correct aberrant splicing caused by pseudoexon sequence inclusion into the mature mRNA. By employing Cas9 fused to a human exonuclease (TREX2), EDSpliCE achieves targeted enhanced deletions of sequences involved in pseudoexon recognition, thereby restoring correct splicing of the pre-mRNA. By addressing two isolated (*ABCA4*:c.5197-557G>T and *USH2A*:c.7595-2144A>G) and two clustered (*ABCA4*:c.5196+1013A>G and *ABCA4*:c.5196+1056A>G) pathogenic deep-intronic variants, we demonstrated effective splicing rescue in minigene assay employing distinct single gRNAs. Further validation in patient-derived fibroblasts for the common *USH2A*:c.7595-2144A>G variant confirmed consistent and high splicing correction. Additionally, the characterization of achieved gene editing affirmed the generation of enhanced deletions by EDSpliCE, revealed high directionality of editing events for all the single gRNAs tested in patient-derived fibroblasts and did not show higher off-target editing potential on selected loci.

**Conclusions:** The successful implementation of the EDSpliCE platform for splicing correction and modulation offers a promising and versatile gene editing approach to address splicing defects, potentially providing a safer option to existing gene editing strategies.

## Background

Correct splicing is essential to guarantee the production of correct mRNA transcripts in eukaryotic cells, fundamental for translation into functional proteins (1). The process is regulated by a complex protein machinery known as spliceosome (2). It recognizes *cis*-acting sequences in the pre-mRNA molecule (i.e. splicing signals), which together orchestrate the recognition of intron-exon boundaries, resulting in the removal of introns from the pre-mRNA molecule. In the last decade, the increasing number of identified disease-causing variants, which affect correct splicing and result in aberrant transcript formation, has underscored the role of missplicing of transcripts as a frequent pathomechanism in inherited disorders (3). Additionally, the unique pathogenic molecular mechanism associated with variants impacting splicing has unveiled new avenues for the exploration of innovative therapeutic strategies, in particular for deep intronic variants inducing pseudoexon activation, which can account for a considerable fraction of disease-linked variants (or alleles) in some genes (4,5,6).

Notably, antisense oligonucleotides (ASOs) and CRISPR/Cas9 genome editing have been harnessed to effectively interfere with aberrant splicing eventually restoring regular splicing (7,8,9). ASOs function through masking *cis*-acting sequences on the pre-mRNA which trigger aberrant splicing, thereby preventing their recognition by the spliceosome. As a result, the formation of correct mature mRNA transcript is restored (10). However, in the context of therapeutic translation, splicing correction achieved by ASOs requires recurrent re-administration of the therapeutic compound and thus incompatible with a single curative treatment (11).

Conversely, the use of CRISPR/Cas9 genome editing potentially enable permanent repair of the splicing defect. Current genome editing approaches to rescue missplicing, involving pseudoexon inclusion due to deep-intronic variants, primarily rely on the use of pairs of single guide RNAs (gRNAs) targeted to generate a genomic deletion encompassing the deep-intronic variant and part, or the whole, pseudoexon sequence (12, 13, 14, 15). Approaches based on single gRNAs designed to target sequences in close proximity with those involved in aberrant splicing have also been demonstrated (14,16). However, although being more minimalistic - thereby preferable for clinical translation - splicing correction induced by single gRNAs is limited in its design by the immediate sequence context and by the fraction of ‘productive’ editing events (small indels) capable of restoring regular splicing (17,18,19). Since the type and fraction of mutational profiles induced by the genome editing hinge on the genomic context as well as cell type, encouraging pre-clinical results obtained in standard cell cultures or advanced cellular models might not be sufficient to deduce comparable results in the *in vivo* scenarios (17,20). Indeed, currently, there is no example of clinical trials for splicing modulation that employs single gRNAs. Conversely, EDIT-101 (NCT03872479) was the first in-human clinical trial aiming to rescue a deep intronic splicing defect in the *CEP290* gene. The approach used a pair of single gRNAs to eliminate the genomic sequence containing the *CEP290*:c.2991+1655A>G deep-intronic variant and corresponding pseudoexon sequence (12). Nonetheless, latest developments are focusing towards editing designs that avoid multiple the generation of double-strand breaks (DSB, e.g. base editing and prime editing) as safer therapeutic options. The main concern regards chromosomal instability following the generation of single or multiple DSB(s), including chromosomal translocations and rearrangements (21,22,23).

With this in mind, we designed a novel CRISPR/Cas9 platform, namely Enhanced-Deletion Splicing Correction Editing (EDSpliCE) (**Figure 1**), to rescue splicing defects induced by pathogenic deep-intronic variants, which lead to pseudoexon inclusion, and validated EDSpliCE exemplarily in variants causing inherited retinal diseases. Initially, four different disease-causing variants were selected for preliminary experiments in minigene assay: the variant c.5197-557G>T, and the clustered variants c.5196+1013A>G and c.5196+1056A>G in *ABCA4* linked to Stargardt disease, the most frequent form of inherited juvenile macular degeneration (or cone-rod dystrophy), and the common c.7595-2144A>G variant in *USH2A* linked to autosomal recessive Usher syndrome or isolated Retinitis pigmentosa (24,25,26,27,28,29,30). Notably, *ABCA4* and *USH2A* are large genes and thus difficult to address therapeutically by means of conventional adeno-associated virus (AAV)-based gene augmentation strategy.

**Figure 1:**
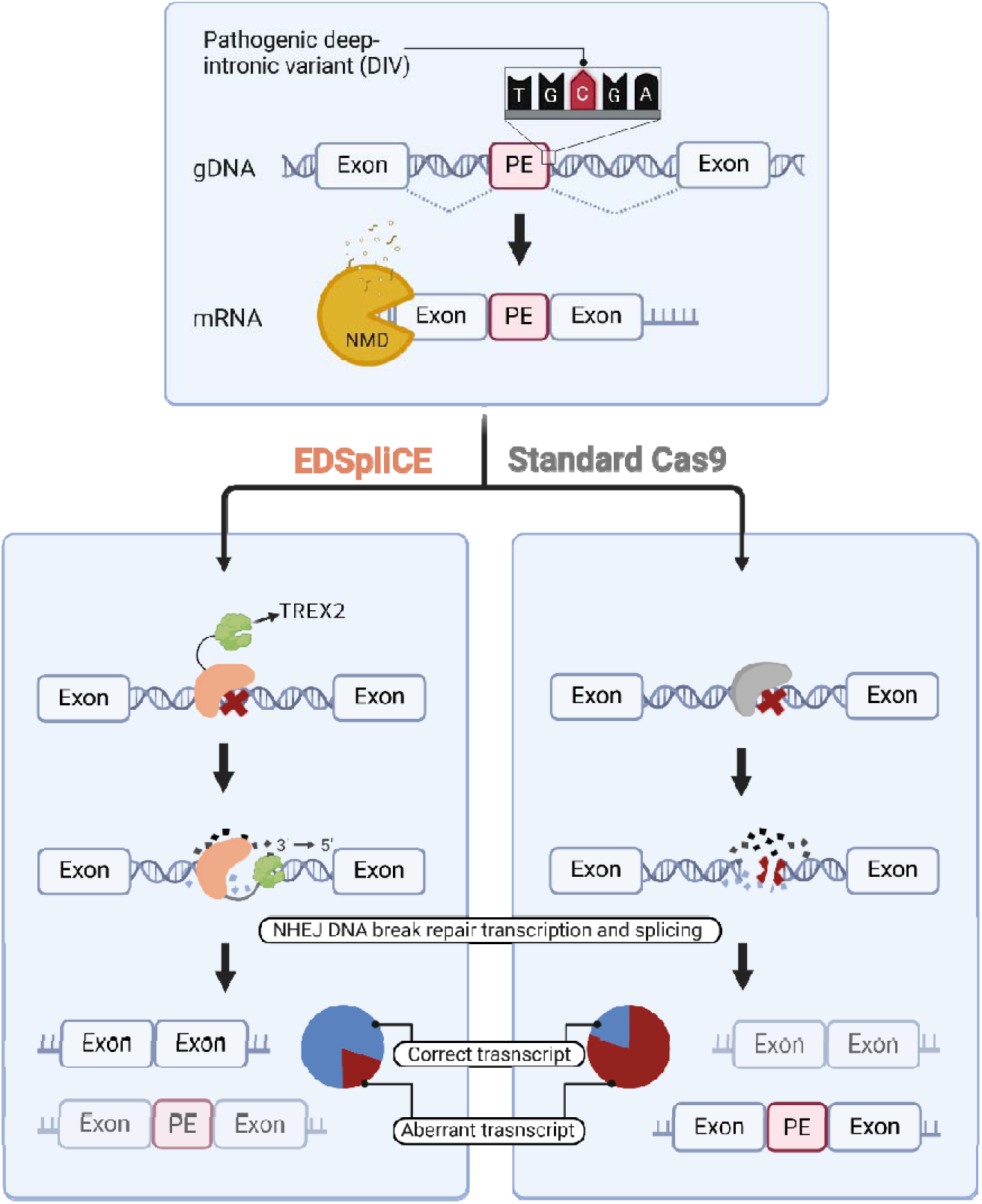
Schematic overview of EDSpliCE vs. standard Cas9 for rescuing splicing defects involving pseudoexon activation. (Top) A pathogenic deep-intronic variant (marked by a red X) induces a splicing defect in which an intronic sequence is erroneously recognized as an exon (pseudoexon, PE), which is included in the mature mRNA transcript. Pseudoexon inclusion frequently results in a frameshift in the open reading frame and premature translation termination, which eventually triggers nonsense-mediated mRNA decay (NMD) pathway. (Bottom) Illustration of the Enhanced-Deletion Splicing Correction Editing (EDSpliCE) platform to rescue splicing defects involving pseudoexon inclusion in comparison to the standard Cas9 approach. Upon Cas9 mediated double strand break formation, the fusion partner, TREX2, further resects the generated 3’ ends, resulting in “enhanced deletions”. In contrast, standard Cas9-mediated disruption of sequences involved in the faulty splicing process primarily relies on the mutational repair profile following the non-homologous end joining (NHEJ) repair mechanism.

The EDSpliCE platform, designed to target aberrant splicing events involving pseudoexon inclusion, implements a chimeric Cas9 molecule, where Cas9 is fused to the human three prime repair exonuclease 2 (TREX2) (17,31). The resulting fusion protein, termed Enhanced-Deletion Cas9 (EDCas9), is coupled to individual single gRNAs and targeted to sequence elements implicated in missplicing (i.e. the cryptic acceptor, donor splice sites, and/or the deep-intronic variant). Owing to the combined activity of Cas9 and TREX2, the occurrence of ‘productive’ deletions is notably increased with EDCas9, leading to the effective perturbation of the targeted DNA sequences, thereby preventing spliceosome recognition of the pseudoexon and thus splicing restoration (17,32). As a mode of action, following a single DSB generated by Cas9, TREX2 processes the resulting DNA ends, thereby promoting the generation of deletions. As opposed to wild-type Cas9 coupled to individual single gRNAs, the generation of EDCas9-mediated deletions is not predominantly determined by the expected repair outcomes (17). This enables more predictable and significant improved splicing rescue regardless of the cell line used. Furthermore, concerning safety, the use of EDCas9 drastically mitigates the generation of chromosomal translocations, thereby enhancing the overall safety profile of the approach to a level comparable to base editors (33,34). To conduct comparison experiments, wild-type Cas9 coupled to the same single gRNAs was used.

## Methods

### gRNA design and off-target prediction

Suitable gRNAs were designed on Benchling.com. Off-target prediction was performed on Off-Spotter (https://cm.jefferson.edu/Off-Spotter/). Based on the resulting list, the three first ranked ones were selected for off-target analysis through high-throughput sequencing.

### Plasmids

The mutant (*USH2A*:c.7595-2144G) and the wild-type (*USH2A*:c.7595-2144A) minigene plasmids were generated by Q5® High-Fidelity DNA Polymerase (New Englands Biolabs) amplification of the target *USH2A* region using gDNA of a human heterozygous *USH2A*:c.7595-2144A/G patient and subsequent cloning into the pSPL3 backbone vector by NEBuilder® HiFi DNA Assembly Cloning Kit (New England Biolabs). The intronic sequence encompassing the *USH2A*:c.7595-2144 location was introduced in the recipient pSPL3 exon trapping vector, including the intronic sequence 966 bp upstream and 914 bp downstream of the *USH2A*:c.7595-2144 location. Primers are listed in **Supplementary Table 5**.

A comparable cloning procedures were used to generate the minigene plasmids for *ABCA4*:c.5197-557G>T, *ABCA4*:c.5196+1013A>G and c.5196+1056A>G. as previously described (14,15).

The standard Cas9 plasmid is represented by PX458 (addgene #48138). To generate the EDCas9 (3xFLAG-SV40 NLS-TREX2-linker-SpCas9) plasmid, the 3xFLAG-SV40 NLS-TREX2 sequence was cloned at the N-terminus of Cas9 by NEBuilder® HiFi DNA Assembly Cloning Kit. Specifically, the PX458 vector was digested by *Age*I and *Eco*RV, the 3xFLAG-SV40 NLS was amplified from the PX458 vector, the TREX2-linker fragment was amplified from the pKLV2.2-TREX2-linker-Cas9 plasmid (generously provided by Dr. Andrew Bassett, Sanger Institute, Hinxton, UK), and the part of the Cas9 sequence digested was amplified back from PX458. Q5® High-Fidelity DNA Polymerase was used for amplification. The fragments and backbone plasmids were cloned following the manufacturer’s protocol. Primers are listed in **Supplementary Table 5**.

The Zhang Lab General Cloning Protocol was used to insert annealed synthetic-oligonucleotide gRNA into the *Bbs*I (New Englands Biolabs) restriction site to clone single gRNAs into PX458 and EDCas9. gRNA sequences are listed in **Supplementary Table 5**.

All the plasmids were prepared endotoxin free using the EndoFree Plasmid Kit (Qiagen) following the manufacturer’s protocol.

Plasmid sequences are provided in **Supplementary Table 6.**

### Sanger sequencing

For sequencing PCR amplification products resulting in multiple bands, the PCR products were cloned by CloneJET PCR Cloning Kit (Thermo Fisher Scientific) following the manufacturer’s protocol. Plasmid constructs were verified by Sanger sequencing using the sequencing primers listed in **Supplementary Table 5**. Plasmid DNA were extracted from bacterial cultures using Monarch Plasmid Miniprep Kit (New England Biolabs) and sequenced using the BigDye Terminator v.1.1 kit (Thermo Fisher Scientific) according to the manufacturer’s protocol, respectively. The same sequencing protocol was used to verify the success of gRNA cloning in the different backbone vectors. Sequencing of PCR amplicons resulting in a single band was carried out using the BigDye Terminator v.1.1 kit according to the manufacturer’s protocol. Sequencing reactions were resolved on an ABI PRISM 3130xl Genetic Analyzer.

### Cell lines and culture conditions

HEK293T cells (ATCC, 293T/17) were cultured in Dulbecco’s modified Eagle’s medium (DMEM; Thermo Fisher Scientific, #41966029) supplemented with 10% fetal bovine serum (FBS; Thermo Fisher Scientific, #10270106), 101 U/mL penicillin/streptomycin (PenStrep; Thermo Fisher Scientific, #15140122) at 37°C in a 5% CO_2_ humidified atmosphere.

A skin biopsy of a *USH2A* patient was obtained upon informed written consent complying with the guidelines and approved by the local ethics committee (Project no. 124/2015BO1). Patient-derived fibroblast cells were expanded and cultured in DMEM supplemented with 20% FBS and 10 1U/mL PenStrep at 37°C in a 5% CO_2_ humidified atmosphere. The genotype of the cell line is: *USH2A*:c.[7595-214A>G];[7595-214A>G] (further denoted as “homozygous” fibroblasts).

### Transfection of cell lines

HEK293T cells were seeded in a 24-well plate (250,000 cells/well) in DMEM without PenStrep, followed by overnight incubation at 37°C in a 5% CO_2_ humidified atmosphere. Cells were transfected using Lipofectamine 3000 (Thermo Fisher Scientific) with 500 ng of total plasmids (copy ratio for minigene assay 1:10—minigene:Cas9 or EDCas9 plasmid). After 24 hours, the cell medium was changed, and cells were harvested 48 hours post-transfection for mRNA isolation.

Homozygous fibroblasts were transfected using the Neon electroporation system, according to the manufacturer’s instructions (Thermo Fisher Scientific). Briefly, cells were detached by Trypsin-EDTA (0.05%) (Thermo Fisher Scientific) (5 min at 37°C), harvested in 10 mL of DMEM and collected by centrifugation at 300 × g for 6 min. To prepare for electroporation, 500,000 cells/reaction were resuspended in 100 μL of Buffer R, and 5 μg of endotoxin-free plasmid (editing plasmid: cells =∼250,000:1) was used per electroporation reaction. Endotoxin-free plasmids were prepared using the EndoFree Plasmid Maxi Kit (QIAGEN) following the manufacturer’s protocol. Electroporation was performed at 1,400 V, 20 ms, 2 pulses. Electroporated cells were immediately plated in a well of a 6-well plate with 2 mL of DMEM without PenStrep.

### Fluorescence-activated cell sorting (FACS)

After 24 hours, electroporated homozygous fibroblasts were sorted for EGFP+ cells. Cells were washed in PBS and detached by Trypsin-EDTA (0.05%) (5 min at 37°C), harvested in 10 mL of DMEM and collected by centrifugation at 300 × g for 6 min. The cell pellet was resuspended in 300 µL PBS. Cell sorting was performed on a MA900 Multi-Application Cell sorter (Sony Biotechnology). The cells were first gated for forward and side scattering, and then EGFP intensity was measured by 488 nm blue laser. Maximal amount of cells was sorted (5,000 to 50,000 cells) and plated back until confluence was reached.

### Splicing analysis

Total RNA of minigene-transfected HEK293T cells and patient-derived fibroblasts was extracted using the peqGOLD Total RNA Kit (VWR Life Science). One microgram of RNA was treated with 1 U of DNaseI (Sigma-Aldrich) following the manufacturer’s instructions for 15 min at room temperature, followed by a 10-min heat inactivation step at 70°C after addition of 1 μL of stop buffer. DNaseI-treated RNA samples were used for cDNA synthesis. The Maxima H Minus First Strand cDNA Synthesis (Thermo Fisher Scientific) was used for HEK293T-derived samples and the SuperScript™ IV First-Strand Synthesis System (Thermo Fisher Scientific) was used for fibroblasts-derived samples. A plasmid-specific primer (pSPL3_SA2_R) was used for cDNA synthesis of HEK293T-derived samples, while random hexamers were used in the case of fibroblasts-derived samples. In both cases the manufacturer’s protocol was followed.

Two microliters of the cDNA were used for PCR amplification employing Taq polymerase (Genaxxon Bioscience) for HEK293T-derived samples and Q5® High-Fidelity DNA Polymerase (New England Biolabs). Primers are listed in **Supplementary Table 5**. PCR products were purified using AMPure XP beads (Beckman Coulter) as per manufacturer’s protocol. Purified samples were analyzed on a 2100 Bioanalyzer instrument employing DNA 1000 Kit reagents (Agilent Technologies) according to the manufacturer’s protocol. The percentage of correctly spliced transcripts (CT) was calculated using the formula: (CP/[CP + AP]) × 100, where CP and AP are the molarity of the fragment corresponding to the correctly spliced RT-PCR product and aberrantly spliced RT-PCR product(s), respectively.

### High-throughput sequencing library preparation for characterization and quantification of editing profiles, as well as off-target assessment

The peqGOLD Tissue DNA Mini Kit (VWR Life Science) was used to extract genomic DNA after genome editing in accordance with the manufacturer’s instructions. A first PCR amplification was used to amplify the target region by Q5® High-Fidelity DNA Polymerase using 10 ng of genomic DNA. To attach Nextera Read adapters to the 5’ end, a second PCR amplification using KAPA HiFi HotStart ReadyMix (Roche), hybrid primers and x35 cycles was performed with 1 µL of 1:500-diluted template of the first amplification. Subsequently, a third round of PCR amplification of 25 cycles was carried out using KAPA HiFi HotStart ReadyMix to add dual indexes and the Illumina i5 and i7 adapters, with the primers listed in **Supplementary Table 5**. The PCR products were then purified with AMPure XP beads following the manufacturer’s protocol, and the purified PCR products were quantified using the AccuBlue NextGen dsDNA Quantitation Kit (Biotium) reagents and a Spark microplate reader (Tecan Life Sciences). The quantified PCR products were then combined in equal amounts to create a library with a final concentration of 10 μM. The library was sequenced at the c.ATG/NGS Competence Center Tübingen core facility of the University Hospital Tübingen on a Miseq. CRISRPesso2 (https://crispresso.pinellolab.partners.org/) was used for alignment, characterization and quantification of the different editing profiles. For the editing efficiency experiments, the results are presented as a percentage of total reads normalized to the mean of the allelic reads of samples treated with a mock gRNA coupled to EDCas9 or Cas9. For off-target experiments, the results are presented as percentage of modified (edited) and non-modified (non-edited) reads.

### Statistics

Statistical analysis was performed on GraphPad Prism (GraphPad Software, La Jolla, CA) using the two-way t-test for data in **Figure 3A**, **Figure 3C**, and **Figure 4**, and **Supplementary Figure 5**, and one-way ANOVA test for data in **Figure 3B**.

**Figure 2:**
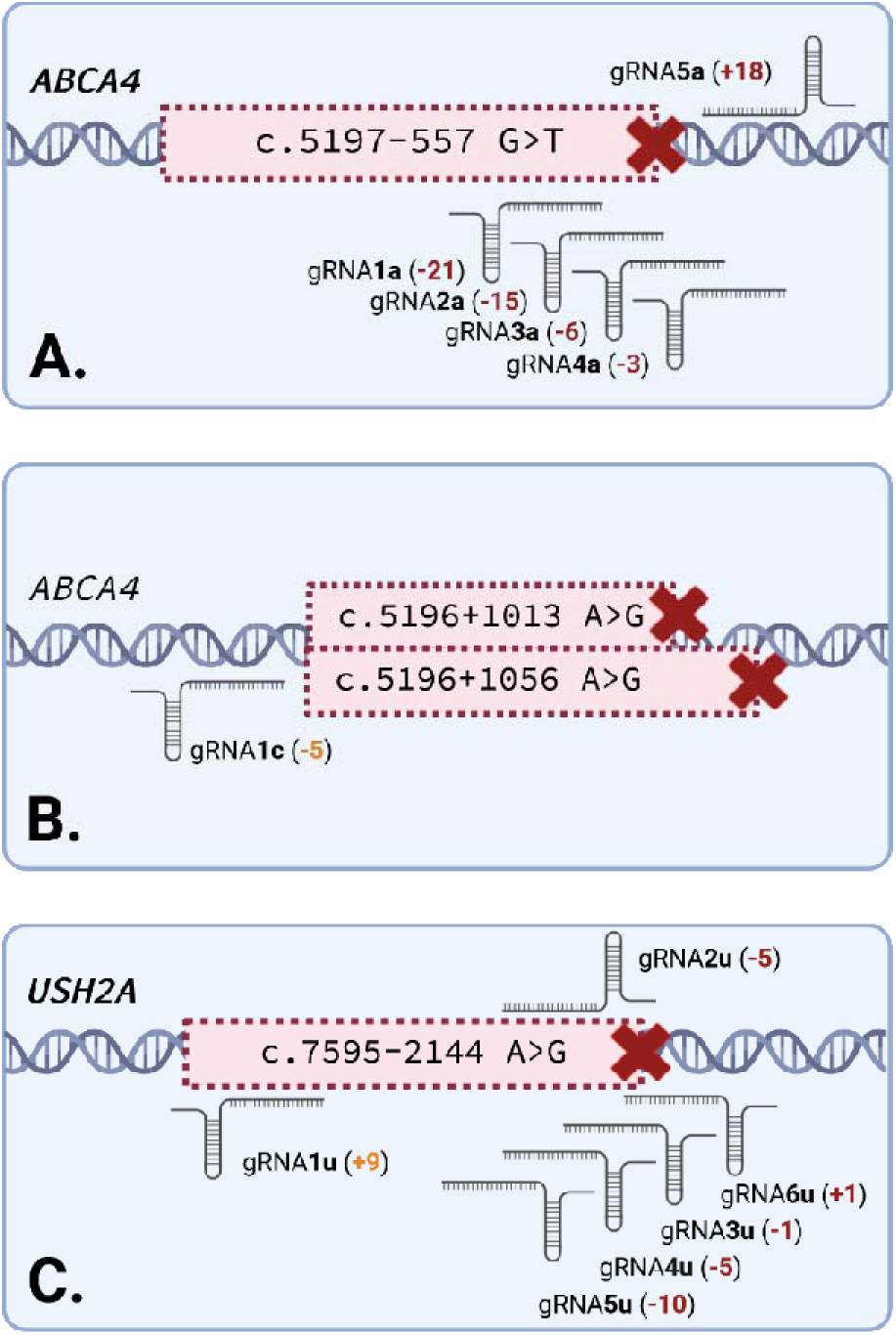
Locations of the single gRNAs used to target the pseudoexons in *ABCA4* and *USH2A*. **(A)** The 188 bp pseudoexon in intron 36 of ABCA4 induced by the isolated c.5197-557G>T deep-intronic variant. **(B)** The 129 bp and 177 bp pseudoexons with shared 5’ acceptor site in intron 36 of ABCA4 induced by the clustered c.5196+1013A>G and ABCA4:c.5196+1056A>G deep-intronic variants, respectively. **(C)** The 152 bp pseudoexon in intron 40 of USH2A induced by the inclusion of the common c.7595-2144A>G deep-intronic variant. **(A,B,C)** In brackets in bold red, the exact position of the gRNA cleavage site in relation to the deep-intronic variant. In brackets in bold orange, the exact position of the gRNA cleavage site in relation to the 5’ acceptor or 3’ donor splice site(s).

**Figure 3:**
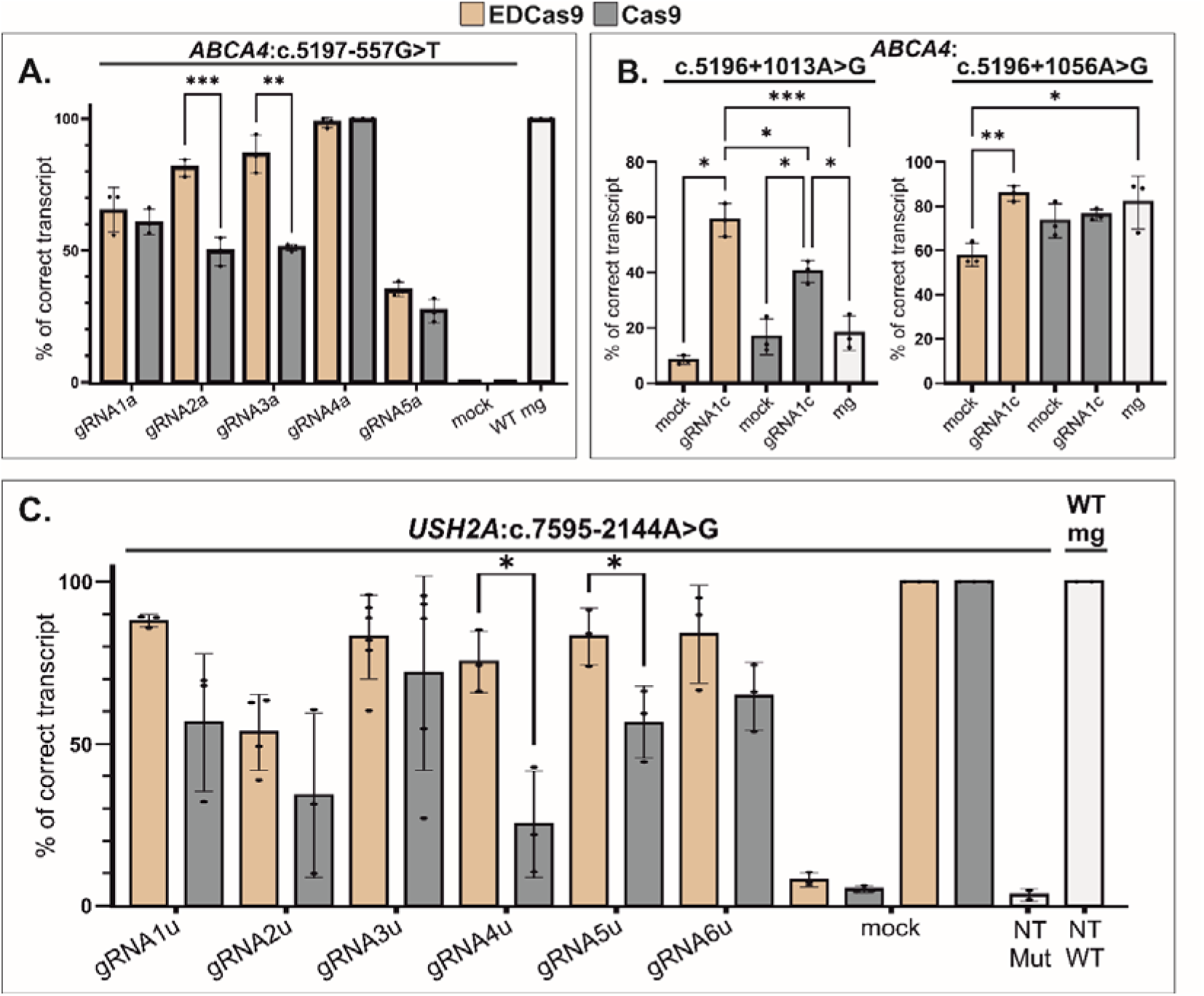
EDCas9- and Cas9-mediated rescue of diverse deep intronic splicing variants on minigene assays in HEK293T. HEK293T cells were co-transfected with mutant (Mut) or wild-type (WT) minigene constructs and plasmids encoding for the different sgRNA/EDCas9 or /Cas9 combinations. **(A)** Bar graphs showing the rescue of the splicing defects induced by the isolated ABCA4:c.5197-557G>T variant, (**B**) the clustered ABCA4:c.5196+1013A>G and c.5196+1056A>G variants, and (**C**) the USH2A:c.7595-2144A>G variant. The relative proportions (percentage) of correctly spliced transcript as quantified from chip automated electrophoresis of RT-PCR products are shown. Co-transfections with a plasmid expressing EDCas9 or Cas9 and a scrambled gRNA were used as controls (mock) as well as sole transfections with the mutant (NT Mut) or the wild-type (NT WT) minigene construct (**Supplementary Figure 1** and **Supplementary Figure 2A-B**). Results are presented as mean ± SD (n = 2–6 independent transfections, single data points are shown, **Supplementary Table 1**). Statistically significant changes in % of correctly spliced transcript are expressed as ∗p ≤0.05, **p≤0.01, ***p≤0.001, and ****p≤0.0001.

**Figure 4:**
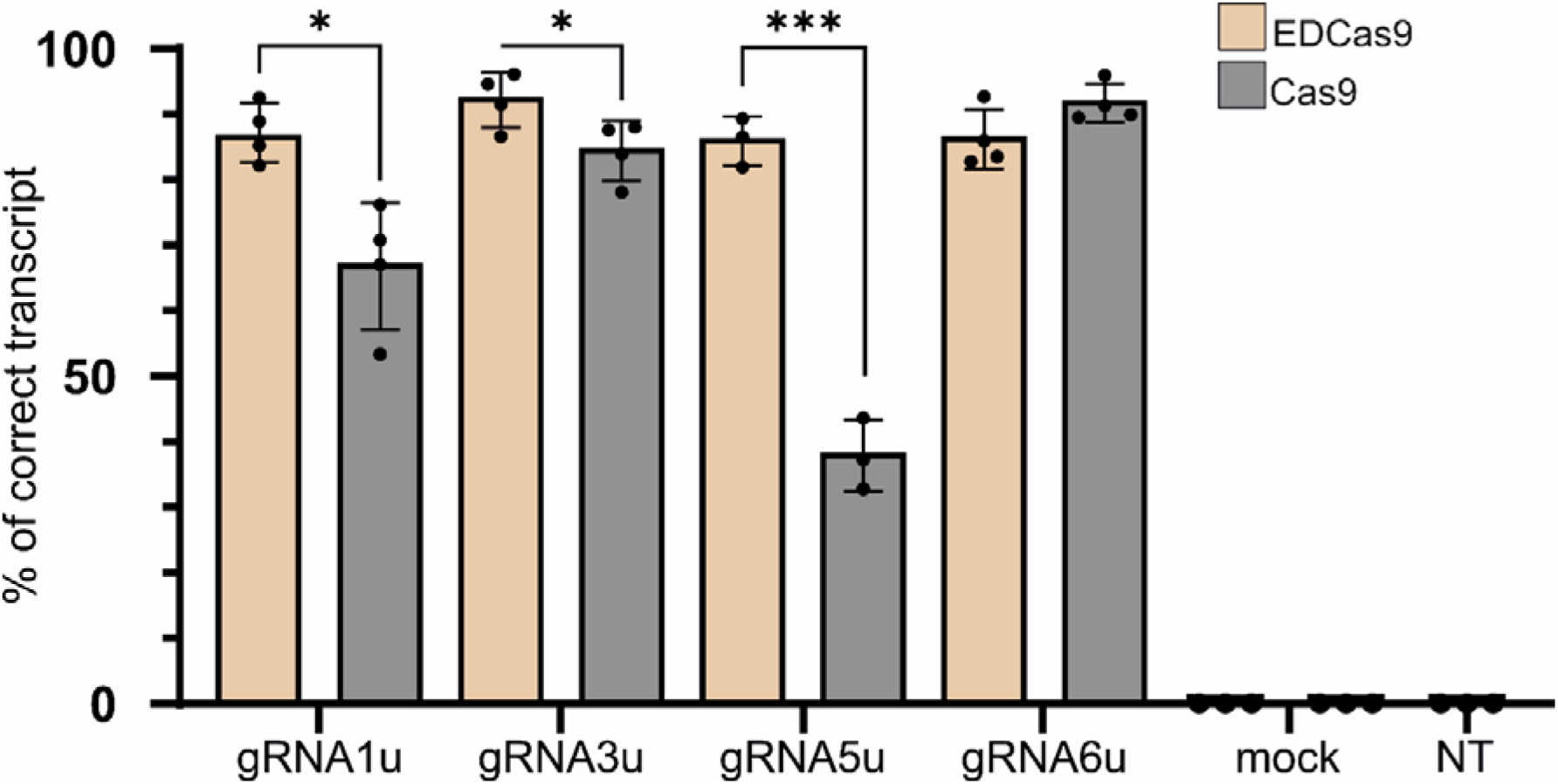
Lead sgRNA/EDCas9 and /Cas9-combinations mediated rescue in homozygous USH2A:c.7595-2144G patient-derived fibroblasts. Patient-derived fibroblasts were electroporated with plasmids encoding for the different lead sgRNA/EDCas9 or /Cas9 combinations and transfected cells enriched by fluorescence-activated cell sorting. Relative proportions (percentage) of correctly spliced transcript as quantified from chip automated electrophoresis of RT-PCR products are shown. Fibroblasts electroporated with a plasmid expressing EDCas9 or Cas9 and a scrambled gRNA were used as a mock gRNA control. Results obtained from non-transfected (NT) fibroblasts are also shown in comparison. Results are presented as mean ± SD (n = 3–4 independent transfections, single data points are shown (**Supplementary Table 1**). Statistically significant changes in USH2A % of correct transcript are expressed as ∗p ≤ 0.05 and ∗∗∗p ≤ 0.001.

## Results

### EDSpliCE improves splicing rescue for the pseudoexon-inducing ABCA4:c.5197-557G>T variant

Genome editing mediated by Cas9 coupled to TREX2 has been shown to increase the occurrence of targeted deletions (17,33,35). We, therefore, speculated that the generation of larger (enhanced) deletions might lead to improved splicing rescue of the missplicing induced by *ABCA4*:c.5197-557G>T variant. The rationale is based on the EDSpliCE-mediated introduction of substantial perturbation of sequences involved in missplicing (i.e. cryptic splice sites and/or deep-intronic variants), which would in turn largely prevent their recognition by the spliceosome, thereby resulting in splicing correction (**Figure 1**).

To test our hypothesis, the *ABCA4*:c.5197-557G>T deep-intronic variant, for which we have previously demonstrated successful single gRNA/CRISPR-Cas9-mediated splicing rescue, was targeted (14). To investigate and compare the efficacy of splicing rescue of the designed single gRNAs in conjunction with EDCas9 (sgRNA/EDCas9) or basic Cas9 (sgRNA/Cas9), we used minigene plasmids which – upon transfection into HEK293T cells - recapitulate the splicing defects induced by the selected variants. Five single gRNAs (gRNA1a, gRNA2a, gRNA3a, gRNA4a, and gRNA5a), that had previously demonstrated splicing rescue efficacy when coupled with SpCas9 (**Figure 2A**), were used (14). These single gRNAs were initially tested in conjunction with two different versions of engineered SpCas9-TREX2 modules: EDCas9 (NLS-TREX2-linker-SpCas9-NLS) and EDCas9-P2A (NLS-SpCas9-P2A-TREX2-NLS). The latter version results in separate TREX2 and SpCas9 polypeptides, while the former produces a chimeric fusion protein with its two domains separated by a flexible linker. Given that preliminary experiments demonstrated superior rescue of EDCas9 over EDCas9-P2A, further experiments were performed using the chimeric EDCas9 fusion protein version (**Supplementary Figure 1**).

Consistent with previous results for SpCas9, HEK293T cells, co-transfected with the mutant *ABCA4*:c.5197-557G>T minigene and a mock gRNA targeting EDCas9 to a protospacer not found in GRCh38, yielded solely misspliced *ABCA4* transcript (14) (**Figure 3A** and **Supplementary Table 1**, **Supplementary Figure 1**). Upon applying the five different single gRNAs (gRNA1a, gRNA2a, gRNA3a, gRNA4a, and gRNA5a) targeting the *ABCA4*:c.5197-557G>T, EDCas9 resulted in a greater fraction of correctly spliced *ABCA4* transcript with four out of five tested sgRNAs compared to Cas9 (**Figure 3A** and **Supplementary Figure 1**), with gRNA2a and gRNA3a showing considerably higher amounts of correct transcript: 86.5±7.3% vs 51.1±1.3% and 81.3±3.3% vs 49.6±5.4%, respectively. For gRNA1a and gRNA5a a slight increase in the fraction of correctly spliced transcript was detected: 64.7±7.8% vs 60.9±4.85% and 35.2±2.8% vs 27.1±4.5%, respectively. Finally, for gRNA4a, both EDCas9 and Cas9 resulted in highly efficient rescue of the splicing defect with only minute (1.5±1.5%) or undetectable levels of misspliced products, respectively.

### EDSpliCE efficiently rescues aberrant splicing for other pseudoexon-inducing deep intronic variants

In order to test its versatility and broad applicability, we then tested EDSpliCE to rescue two *ABCA4* clustered deep-intronic variants (c.5196+1013A>G and c.5196+1056A>G) in intron 36 and the common c.7595-2144A>G variants in intron 40 of *USH2A*, again benchmarked against standard Cas9.

Both additional *ABCA4* variants (c.5196+1013A>G and c.5196+1056A>G) induce splicing defects resulting in partial missplicing of the minigene-derived transcripts in HEK293T cells (15). Notably, co-transfection of the minigene constructs together with the Cas9 construct expressing a scrambled gRNA (mock) consistently yielded a higher proportion of correctly splicing products compared to EDCas9 mock, thereby setting more elevated thresholds for splicing correction (**Figure 3B**, **Supplementary Table 1**, and **Supplementary Figure 2A**). Irrespective, gRNA1c co-expressed with EDCas9 resulted in higher levels of splicing rescue compared to its co-expression with Cas9 (**Figure 3B**). Specifically, gRNA1c/EDCas9 yielded 56.0±4.9% and 85.6±2.9% of correctly spliced *ABCA4* transcripts for c.5196+1013A>G and c.5196+1056A>G, respectively, whereas gRNA1/Cas9 achieved 40.3±3.3% and 76.0±2.2% (**Figure 3B**, **Supplementary Table 1**, and **Supplementary Figure 2A**). If the fold-change to the thresholds of correct *ABCA4* transcript set by mock-transfected samples is used for normalization (**Supplementary Table 2**), gRNA1c/EDCas9 showed a 580±60% increase for c.5196+1013A>G and a 50±10% increase for c.5196+1056A>G of correct *ABCA4* transcript compared to 140±20% and 0±10% induced by gRNA1c/Cas9 for the same deep-intronic variants, respectively.

For the *USH2A*:c.7595-2144A>G variant, we first validated our minigene construct and minigene splicing assay in transiently transfected HEK293T cells (**Supplementary Figure 3**). Transfection of the mutant and wild-type minigene constructs resulted in 3.4±2.0% and 100±0% correct *USH2A* transcript, respectively (**Figure 3C, Supplementary Figure 2B** and **Supplementary Table 1**). Co-transfection of the mutant or the wildtype minigene with either EDCas9/mock-gRNA or Cas9/mock-gRNA (protospacer absent in GRCh38), demonstrated negligible effects on minigene splicing (7.9±2.2% and 5.1%±1.0% correctly spliced *USH2A* transcripts for the mutant minigene, and 100±0% correctly spliced *USH2A* transcripts for the wildtype minigene). Next, we tested a series of single gRNA targeting the c.7595-2144A>G – induced pseudoexon in combination with EDCas9 or Cas9 to rescue the splicing defect. Overall, all tested sgRNA/EDCas9 combinations performed superior in splicing rescue compared to the corresponding sgRNA/Cas9 combinations (**Figure 3C, Supplementary Figure 2B** and **Supplementary Table 1**). With EDCas9, the highest efficacy peaked at 88.0±1.9% of correctly spliced transcripts with gRNA1u, and five of the six tested gRNAs showed levels of correctly spliced transcripts of 70% or higher. In contrast, Cas9 reached a maximum efficacy of 71.8±30.0% of correctly spliced transcripts with gRNA3u and only one of the six tested gRNAs reached a level of 70%.

### EDSpliCE induces high and consistent splicing rescue in patient-derived homozygous USH2A:c.7595-2144G fibroblasts

Given its therapeutic relevance due to its high allele frequency (**Supplementary Table 3**), the pathogenic *USH2A*:c.7595-2144A>G deep-intronic variant was prioritized for subsequent experiments in patient-derived cells. gRNA1u, gRNA3u, gRNA5u, and gRNA6u, showing the highest efficacy in rescuing the splicing defect in the minigene assay in HEK293T cells, were chosen for further validation in homozygous *USH2A*:c.7595-2144G patient-derived fibroblasts. Electroporation was used to deliver the genome editing plasmids into the cells. To limit the variability across electroporation experiments, we used EDCas9 or Cas9 constructs with in-frame expressed EGFP, and sorting of EGFP-positive cells 24 hours post electroporation. Sorted cells were sub-cultured until confluent. To prevent possible degradation of the aberrant *USH2A* transcript by nonsense-mediated mRNA decay (NMD), fibroblasts were treated with 0.1 mg/mL cycloheximide (CHX), a commonly used NMD blocker (36), 16 hours prior to harvesting.

Mock-transfected *USH2A*:c.7595-2144G fibroblasts showed a fully penetrant splicing defect with not even marginal levels of the correctly spliced transcript detectable (**Figure 4, Supplementary Figure 4A**). All four tested sgRNA/EDCas9 combinations resulted in high levels of correctly spliced transcript, ranging from 85.7±3.7% for gRNA5u to 92.4±4.8% for gRNA3u (**Supplementary Table 1, Figure 4**). These results are in line with the results obtained with the minigene assay in HEK293T cells. A less consistent increase in the fraction of correct *USH2A* transcripts (range: 38.2±5.4% for gRNA5u to 91.8±3.2% for gRNA6u) was also observed with Cas9. Yet, as opposed to EDCas9 results, there were notable differences in splicing rescue induced by Cas9 between the minigene assay in HEK293T cells and patient-derived fibroblasts, with gRNA5u (56.7±11.2% and 38.2±5.4%, respectively) and gRNA6u (64.7±10.4% and 91.8±3.2%, respectively) showing the greatest variance between the two experimental model systems.

Using the EDCas9 fusion, gRNA1u, gRNA3u, and gRNA5u resulted in significant higher fraction of correctly spliced transcript compared to Cas9 (**Figure 4**). Specifically, gRNA1u resulted in 86.7±5.2% and 66.8±8.7% correctly spliced *USH2A* transcript, gRNA3u in 92.4±4.8% and 84.6±5.0% correctly spliced *USH2A* transcript, and gRNA5u in 85.7±3.7% and 38.2±5.4% correctly spliced *USH2A* transcript, respectively. In contrast, gRNA6u showed comparable splicing rescue efficacies for both editing enzymes (86.3±4.9% and 91.8±3.2%, respectively).

To further characterize editing induced outcomes at the transcript level, residual aberrant *USH2A* transcripts from the treated fibroblasts were subjected to sub-cloning and sequencing and revealed the presence of further missplicing products (**Supplementary Figure 4B**). In detail, we observed shorter pseudoexons of 134 bp and 136 lacking a portion of the 5’ end of the pseudoexon in the case of gRNA1/EDCas9. Additionally, the characterization of the residual misspliced transcripts of gRNA5/EDCas9 showed inclusion of pseudoexons lacking 3 bp or 4 bp at the 3’-terminal end, while gRNA5/Cas9 retained pseudoexons lacking 1 bp, 2 bp, or 4 bp. Notably, gRNA5/Cas9 also retained pseudoexons with single or double nucleotide insertions. In the cases of gRNA1/Cas9 and gRNA6/Cas9, “hybrid” pseudoexons were detected, where part of the retained sequence was derived from another gene, which might indicate an off-target site mediating this chromosomal translocation. Specifically, the sequence NG_050857.1:85744-85719 was detected for gRNA1/Cas9 and NG_0047091.1:51092-51136 was detected for gRNA6/Cas9, respectively.

### EDSpliCE induces larger and directional deletions while exhibiting comparable off-target profile on selected loci

To quantify and profile the genomic DNA cleavage activity of EDCas9 and Cas9 coupled to the lead single gRNAs targeting *USH2A*:c.7595-2144G, genomic DNA of the treated *USH2A*:c.7595-2144G fibroblasts was subjected to targeted high-throughput sequencing (HTS). A genomic region of 425 bp for gRNA1u- and 439 bp for gRNA3u-, gRNA5u-, and gRNA6u-treated fibroblasts, with amplicons covering the respective gRNA target sequence, underwent single-read sequencing (x500 cycles). The analysis of the data showed comparable genomic DNA editing activity for EDCas9 and Cas9, as defined by the percentage of indel reads (i.e. deletions or insertions) (**Supplementary Figure 5**). However, the shape of the resulting deletion profiles varied substantially. While Cas9 treatment predominantly resulted in small indels of ≤5 nucleotides (gRNA1: 61.2±1.9%; gRNA3: 69.8±0.8%; gRNA5: 79.1±3.7%; gRNA6: 76.8±7.3%), EDCas9 induced consistent and directional deletion of larger sequence stretches (**Figure 5**). Across the lead gRNAs, EDCas9 treatment resulted in an average of 92.3±3.2%, 62.6±15.3%, and 12.1±5.3% of deletions of ≥5, ≥15, and ≥30 base pairs, respectively, while for Cas9 treated fibroblasts these fractions were reduced to 41.3±7.6%, 8.2±4.8%, and 2.2±2.5%, respectively (**Figure 5**). Notably, for EDCas9 we observed a directionality of deletions towards the 3’-end of the non-target strand (**Supplementary Figure 6**). Moreover, insertions were rarely observed with EDCas9. The highest detected fraction of insertions for EDCas9-treated fibroblasts was 0.32±0.32% (2 bp insertion, gRNA1u/EDCas9) compared to 12.9±9.1% for Cas9 (1 bp insertion, gRNA5u/Cas9) (**Supplementary Table 4**). Quantification of the editing efficiency showed gRNA5u having the highest efficacy of 88.2±0.9% and 91.5±1.6% for EDCas9 and Cas9, respectively. Editing with gRNA1u, gRNA3u, and gRNA6u resulted in similar editing outcome levels ranging from 79.1±8.9% for gRNA6u/EDCas9 to 86.1±8.0% for gRNA1u/EDCas9.

**Figure 5:**
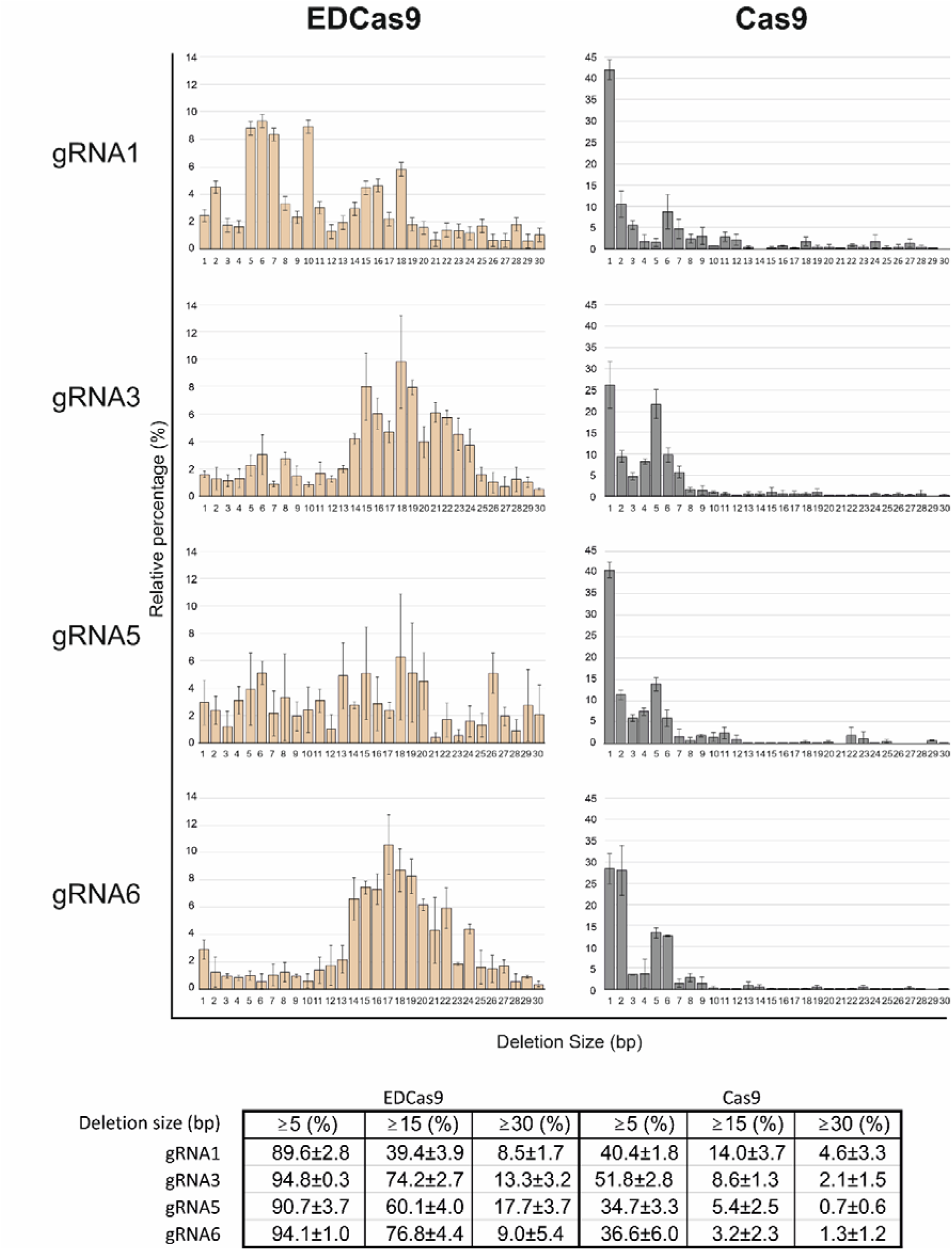
Deletion profiles induced by EDCas9 and Cas9 with lead single gRNAs in homozygous USH2A:c.7595-2144G patient-derived fibroblasts. The X-axis of the plot represents the deletion size in base pairs (bp), ranging from 1 to 30 bp, while the relative percentage (%) of each deletion size is shown on the Y-axis. In the table below, the relative quantification (%) of ≥5, ≥15, and ≥30 bp deletions is given for the EDCas9/gRNA and Cas9/gRNA combinations. Results are presented as mean ± SD (n = 3 independent replicates).

Furthermore, HTS analysis of three off-target sites for gRNA3u and gRNA6u, predicted based on potentially permissive nucleotide mismatches to the gRNA sequence, showed no significant editing differences between EDCas9 and Cas9 (**Supplementary Figure 7** and **Supplementary Figure 8**).

## Discussion

Due to their location remote from coding sequences, increasing therapeutic interest embraces pathogenic deep-intronic variants which affect pre-mRNA splicing. In this respect, antisense oligonucleotides have emerged as effective molecules for splicing correction (7,8,9,11). However, ASOs undergo degradation and, therefore, their effect is transient, rendering recurrent administration necessary. In case of ocular or retinal disease, this goes along with risks of bleeding, cataract and intraocular infection (11,37,38,39). In contrast, by targeting the genomic level, genome editing technologies provide a promising avenue for a curative treatment of splicing defects induced by pathogenic deep-intronic variants (12,13,14,15,16,40). However, current gene editing approaches aiming to treat pseudoexon inclusion-linked splicing defects are typically tailored to induce a deletion of a genomic fragment encompassing the entire pseudoexon sequence, which i.) require two single gRNAs, ii.) have higher risk of chromosomal instability and activation of P53-related pathway, and iii.) have, arguably, higher chance of off- and on-target sequence aberration (14,15,41,42). Alternatively, a single gRNA approach can be used, which relies on perturbation of crucial missplicing sequences (e.g. cryptic splice sites) through mutational outcomes induced by non-homologous end joining (NHEJ) (14,16). Nevertheless, also this approach faces limitations, as only a restricted number of single gRNAs targeting the immediate proximity of the crucial sequences prove effective in substantial splicing rescue when using standard Cas variants (14,16). Moreover, the considerable variability in the resulting indel formations, affecting splicing rescue effectiveness and depending on cell type, poses a significant challenge in scaling this editing approach for clinical applications (17,20).

Given these challenges, we hypothesized that fusing a 3’-end-processing enzyme (TREX2) to Cas9 could increase the frequency and size of deletions, thereby boosting sequence perturbation, which would lead to higher fraction of molecules in which aberrant splicing is rescued (17). Consequently, while necessitating only a single gRNA, the described editing approach not only increases the likelihood of identifying effective single gRNAs but, by expanding the targetable sequence window, also enhances the potential for pinpointing specific ones suitable for clinical applications. Indeed, for the four deep intronic mutations tested in this study, 12 out of 13 gRNAs/EDCas9 combinations resulted in a higher fraction of correctly spliced transcripts compared to when used with Cas9. Of course, there will be (lucky) instances in which Cas9 will suffice, such as gRNA4a which achieved already near complete rescue of the *ABCA4*:c.5197-557G>T-induced splicing defect, most likely due to its cleavage site being close (−3 nt) to the *ABCA4*:c.5197-557G>T variant itself. Yet, EDSpliCE has the potential to considerably expand the spectrum of deep intronic mutations to be efficiently targeted. Remarkably, in the minigene assay experiments, the relative fraction of correct transcript for the clustered pathogenic *ABCA4* splicing defects induced by *ABCA4*:c.5196+1013A>G and *ABCA4*:c.5196+1056A>G was significantly increased by the use of a single gRNAs, highlighting the potential of EDCas9 as an appealing option for targeting splicing defects arising from close pathogenetic deep-intronic variants. It is worth noting that, the variability of correctly spliced *ABCA4* transcript observed for the clustered c.5196+1056A>G deep-intronic variant between EDCas9-mock- and Cas9-mock- and minigene-transfected samples may be attributed to the simultaneous overexpression of two different plasmids (minigene and editing plasmid), which may influence splicing patterns by differently perturbating the transcriptome and altering the levels of NMD-related proteins and/or splicing factors. Such a finding was also recently reported by Suárez-Herrera and co-workers when addressing some *ABCA4* splicing defect in midigene-transfected HEK293 (43).

In patient-derived fibroblasts, subsequent validation experiments of four lead gRNAs (gRNA1u, 3u, 5u and 6u) targeting the common *USH2A:* c.7595-2144A>G splicing defect affirmed robust splicing rescue (>80%) with EDSpliCE, closely resembling the results obtained in the minigene experiments. Notably, differences emerged for Cas9, highlighting the potential impact of diverse mutational profiles in different cell lines on the outcomes of splicing rescue. This was particular evident for gRNA5u, inducing 56.7±11.2% vs 38.2±5.4% of correctly spliced transcript and gRNA6u, resulting in 64.7±10.4% vs 91.8±3.2%, in minigene assay and patient-derived fibroblasts, respectively. Furthermore, in patient-derived fibroblasts, only two single gRNAs applied with Cas9 (gRNA3u and gRNA6u) led to a considerable fraction of >80% correct transcript, limiting the number of effective single gRNAs to two as opposed to four for EDCas9.

It is crucial to highlight that the higher splicing rescue achieved with EDCas9 cannot be attributed to a higher cut efficiency at the genomic level, as this was comparable to that achieved with Cas9 (**Supplementary Figure 5**). Moreover, no differences across three predicted off-target sites for gRNA3u and gRNA6u was detected upon editing with either EDCas9 or Cas9, suggesting a similar off-target activity of these two molecules most likely govern by the type of endonuclease used (i.e. SpCas9) (**Supplementary Figure 7** and **Supplementary Figure 8**). When analyzing the resulting genomic editing profiles, it is evident that EDCas9 boosts the occurrence of deletions as well as the size of these deletions, while drastically reducing the generation of insertions (**Figure 5, Supplementary Table 4**). Most interestingly, the cleavage kinetic of EDCas9 induces highly directional deletions biased towards the 3’-end of the non-target strand (**Supplementary Figure 6**). Specifically, for the tested four single gRNAs, no deletion >1bp was detected in the 5’-end direction of the non-target strand, making EDCas9 highly attractive for gene editing applications requiring directionality (e.g. targeting exon/intron boundaries or inducing targeted large deletions). This characteristic is most likely explained by the preferential release of the 3’-end of the non-target strand upon cleavage, while maintaining stable engagement with the remaining three strands. In this scenario, the released end (3’-end of the non-target strand) is immediately accessible for processing by TREX2, thereby resulting in deletions biased towards this specific direction (44). The deletion distributions induced by EDCas9 for the four tested gRNAs do not appear to follow a single pattern (**Figure 4**), suggesting involvement of the sequence context in determining the final repair outcomes.

In contrast, the editing profile of gRNA6u/SpCas9 predominantly results in the deletion of the deep-intronic variant itself (positioned at −1 relative to the cut site) (**Supplementary Figure 6**). This deletion, in turn, contributes to a remarkably high rescue rate of 91.8±3.2% in patient derived fibroblasts. Conversely, owing to the directional bias of EDCas9, the diverse deletion profile induced by gRNA6u in conjunction with EDCas9 ensure the preservation of the deep-intronic variant, accompanied by extensive perturbation in downstream sequences. Nevertheless, even in this case, the achieved level of splicing rescue remains high (86.3±4.9%).

A crucial aspect of the proposed efficient editing strategy for splicing modulation lies in the use of the human TREX2 exonuclease. It is important to consider that, by definition, TREX2 may possess the capacity to process not only the intended target sequence specified by the gRNA/Cas9 complex but also other free DNA ends, potentially raising safety concerns. However, studies have demonstrated that the inactivation of TREX2’s DNA-binding domain through targeted mutagenesis preserves the catalytic ability of the Cas9-exonuclease (Cas9-exo) fusion endonuclease while eliminating its capacity to bind DNA (31,33,45). This may provide an attractive solution to effectively mitigate safety concerns that might be linked to the proposed editing approach. Conversely, it has been shown that the use of Cas9-exo molecules remarkably improves the safety profile by drastically reducing the occurrence of chromosomal translocations, also in postmitotic cells (45,46). This is also supported by our results on the characterization of residual misspliced products upon editing, which suggests the presence of translocations induced by gRNA1u and gRNA6u in the Cas9 setting, contrasting with their absence when Cas9 is replaced by EDCas9. In general, the suggested molecular mechanism underlying the diminished chromosomal instability mediated by Cas9-exo molecules, which include EDCas9, revolves around averting repetitive cycles of DNA cutting and rejoining. Specifically, upon the cleavage of the target sequence mediated by Cas9, TREX2 promptly processes the resulting DNA ends, thereby preventing NHEJ from reconstituting the target sequence, recurrent gRNA-mediated binding and repeated re-initiation of the cleavage cycle (35). Based on this mechanism, EDCas9 might also result in reduced activation of p53-related pathways triggered upon looping DNA cutting, which, consequently, has the potential to further bolster its safety profile (41,42).

In considering the potential future applications of EDSpliCE for the treatment of inherited retinal dystrophies, delivery of the editing system through AAV particles represents the currently preferred choice (47). Nevertheless, the size of the EDCas9 molecule, coupled with a gRNA cassette, exceeds the AAV cargo capacity, necessitating the implementation of a dual AAV system (e.g. trans-splicing design) (48). Alternatively, the engineering of smaller Cas orthologues (e.g. SaCas9 or Nme2Cas9) could provide a viable solution, as their sizes would fit the capacity of a single AAV plasmid (49,50).

## Conclusions

In conclusion, by demonstrating substantial correction of different splicing defects due to pseudoexon inclusion in minigene assay, and by further confirming the robust correction of the recurrent *USH2A*-related splicing defect in patient-derived fibroblasts, this study shows the potential of the EDSpliCE platform in effectively modulating splicing and encourages its further exploration as a promising alternative to existing editing approaches. Moreover, owing to its remarkable deletion directionality, EDSpliCE represents also a powerful tool for effectively addressing splicing defects resulting from variants located in close proximity to exon/intron boundaries. Ultimately, the versatility of splicing modulation mediated by EDSpliCE opens avenues for purposefully induction of aberrant splicing, which might prove useful in addressing dominant-acting pathogenic alleles.

## List of abbreviations

AAV: adeno-associated virus
ASO: antisense oligonucleotide
Cas9-exo: Cas9-exonuclease CHX: cycloheximide
DSB: double-strand break
EDCas9: Enhanced-Deletion Cas9
EDSpliCE: Enhanced-Deletion Splicing Correction Editing
FACS: Flourescence-activated cell sorting
gRNA: guide RNA
HTS: high-throughput sequencing
NHEJ: non-homologous end joining
NMD: nonsense-mediated mRNA decay
PE: pseudoexon
TREX2: three prime repair exonuclease 2

## Declarations

### Ethics approval and consent to participate

The study was conducted in accordance with the ethical principles for research involving human subjects outlined in the Declaration of Helsinki. The Fibroblast cell line was expanded from a skin biopsy collected from an adult patient upon written informed consent approved by the ethics committee of the Medical Faculty of the University of Tübingen (Project no. 124/2015BO1).

### Consent for publication

Not applicable

### Availability of data and materials

The datasets used and/or analyzed during the current study are available from the corresponding author on reasonable request.

### Authors’ contributions

Conceptualization: PDA; Methodology: PDA, AFT; Investigation: PDA, AFT, SSH, SSP, ER; Resources: KS, LK; Writing - Original Draft: PDA; Writing - Review & Editing: SK, BW; Supervision: PDA, SK, BW; Funding acquisition: SK, PDA.

## Acknowledgements

We thank Dr. Andrew Bassett (Wellcome Trust Sanger Institute, UK) for kindly providing the original pKLV2.2-Cas9-2A-TREX2 plasmid.

