## Supplementary for "EDSpliCE, a CRISPR-Cas9 gene editing platform to rescue splicing, effectively corrects inherited retinal dystrophy-associated splicing defects"

**Supplementary Material**

**Supplementary Table 1:** Percentage (%) values of correctly spliced transcript achieved in minigene (mg) assay and homozygous USH2A:c.7595-2144G patient-derived fibroblasts. Results are presented as mean ± SD. The number of biological replicates performed is indicated.

| **Sample** | **Percentage of correct *USH2A* transcript (%)** | **Number of biological replicates** |
| --- | --- | --- |
| ***Minigene*** | | |
| EDCas9_gRNA1a_MgABCA4:c.5197-557G>T | 64.7±7.8% | 3 |
| EDCas9_gRNA2a_MgABCA4:c.5197-557G>T | 81.3±3.3% | 3 |
| EDCas9_gRNA3a_MgABCA4:c.5197-557G>T | 86.5±7.3% | 3 |
| EDCas9_gRNA4a_MgABCA4:c.5197-557G>T | 98.5±1.5% | 3 |
| EDCas9_gRNA5a_MgABCA4:c.5197-557G>T | 35.2±2.8% | 3 |
| Cas9_gRNA1a_MgABCA4:c.5197-557G>T | 60.9±4.9% | 3 |
| Cas9_gRNA2a_MgABCA4:c.5197-557G>T | 49.6±5.4% | 3 |
| Cas9_gRNA3a_MgABCA4:c.5197-557G>T | 51.1±1.3% | 3 |
| Cas9_gRNA4a_MgABCA4:c.5197-557G>T | 100.0±0.0% | 3 |
| Cas9_gRNA5a_MgABCA4:c.5197-557G>T | 27.1±4.5% | 3 |
| EDCas9-mockgRNA_MgABCA4:c.5197-557G>T | 0.0±0.0% | 3 |
| Cas9-mockgRNA_MgABCA4:c.5197-557G>T | 0.0±0.0% | 3 |
| WT-ABCA4-minigene | 100.0±0.0% | 3 |
| EDCas9_gRNA1c_MgABCA4:c.5196+1013A>G | 56.0±4.9% | 3 |
| EDCas9_gRNA1c_MgABCA4:c.5196+1056A>G | 85.6±2.9% | 3 |
| Cas9_gRNA1c_MgABCA4:c.5196+1013A>G | 40.3±3.3% | 3 |
| Cas9_gRNA1c_MgABCA4:c.5196+1056A>G | 76.0±2.2% | 3 |
| EDCas9_mockgRNA_MgABCA4:c.5196+1013A>G | 8.3±1.2% | 3 |
| EDCas9_mockgRNA_MgABCA4:c.5196+1056A>G | 58.0±4.2% | 3 |
| Cas9_mockgRNA_MgABCA4:c.5196+1013A>G | 16.6±5.2% | 3 |
| Cas9_mockgRNA_MgABCA4:c.5196+1056A>G | 73.3±6.3% | 3 |
| MgABCA4:c.5196+1013A>G | 18.1±5.4% | 3 |
| MgABCA4:c.5196+1056A>G | 81.9±9.5% | 3 |
| EDCas9_gRNA1u_MgUSH2A:c.7595-2144A>G | 88.0±1.9% | 3 |
| EDCas9_gRNA2u_MgUSH2A:c.7595-2144A>G | 53.6±11.8% | 4 |
| EDCas9_gRNA3u_MgUSH2A:c.7595-2144A>G | 83.0±12.8% | 6 |
| EDCas9_gRNA4u_MgUSH2A:c.7595-2144A>G | 75.3±9.6% | 3 |
| EDCas9_gRNA5u_MgUSH2A:c.7595-2144A>G | 83.1±8.7% | 3 |
| EDCas9_gRNA6u_MgUSH2A:c.7595-2144A>G | 83.8±15.2% | 3 |
| Cas9_gRNA1u_MgUSH2A:c.7595-2144A>G | 56.6±11.8% | 3 |
| Cas9_gRNA2u_MgUSH2A:c.7595-2144A>G | 34.0±25.3% | 3 |
| Cas9_gRNA3u_MgUSH2A:c.7595-2144A>G | 71.8±30.0% | 5 |
| Cas9_gRNA4u_MgUSH2A:c.7595-2144A>G | 25.1±16.4% | 3 |
| Cas9_gRNA5u_MgUSH2A:c.7595-2144A>G | 56.7±11.2% | 3 |
| Cas9_gRNA6u_MgUSH2A:c.7595-2144A>G | 64.7±10.4% | 3 |
| EDCas9_mockgRNA_MgUSH2A:c.7595-2144A>G | 7.9±2.2% | 2 |
| Cas9_mockgRNA_MgUSH2A:c.7595-2144A>G | 5.1±1.0% | 3 |
| EDCas9_mockgRNA_WT-USH2A-minigene | 100%±0.0% | 3 |
| Cas9_mockgRNA_WT-USH2A-minigene | 100%±0.0% | 3 |
| MgUSH2A:c.7595-2144A>G | 3.3±2.0% | 2 |
| WT-USH2A-minigene | 100%±0.0% | 2 |
| ***Patient-derived homozygous USH2A:c.7595-2144A>G fibroblasts*** | | |
| EDCas9_gRNA1_Fibroblasts | 86.6±5.2% | 4 |
| EDCas9_gRNA3_Fibroblasts | 92.4±4.8% | 4 |
| EDCas9_gRNA5_Fibroblasts | 85.7±3.7% | 3 |
| EDCas9_gRNA6_Fibroblasts | 86.3±4.9% | 4 |
| Cas9_gRNA1_Fibroblasts | 66.8±8.7% | 4 |
| Cas9_gRNA3_Fibroblasts | 84.6±5.0% | 4 |
| Cas9_gRNA5_Fibroblasts | 38.2±5.4% | 3 |
| Cas9_gRNA6_Fibroblasts | 91.8±3.2% | 4 |
| Cas9_mockgRNA_Fibroblasts | 0.0±.0.% | 3 |
| EDCas9_mockgRNA_Fibroblasts | 0.0±.0.% | 3 |
| Fibroblasts | 0.0±.0.% | 2 |

**Supplementary Table 2:** Calculated percentage (%) of increase in the fraction of correctly spliced ABCA4 transcripts in minigene assay normalized to mock-transfected samples for the selected clustered deep-intronic variants ABCA4:c.5196+1013A>G and ABCA4:c.5196+1056A>G.

| **Sample** | **% increase change vs Mock (%)** |
| --- | --- |
| EDCas9_gRNA1c_MgABCA4:c.5196+1013A>G | 580±60% |
| EDCas9_gRNA1c_MgABCA4:c.5196+1056A>G | 50±10% |
| Cas9_gRNA1c_MgABCA4:c.5196+1013A>G | 140±20% |
| Cas9_gRNA1c_MgABCA4:c.5196+1056A>G | 0±10% |

**Supplementary Table 3: Frequency of the most relevant and common pathogenic deep-intronic variants in the inherited retinal dystrophy field**

| **Deep-intronic variant** | **dbSNP** | **Allele frequency** |
| --- | --- | --- |
| ***CEP290*:c.2991+1655A>G** | rs281865192 | C=0.000295 (78/264690, TOPMED)  **C=0.000286 (40/140018, gnomAD)**  C=0.00021 (4/19368, ALFA) |
| ***USH2A*:c.7595-2144A>G** | rs786200928 | C=0.000068 (18/264690, TOPMED)  **C=0.000043 (6/140282, gnomAD)**  C=0.00021 (3/14050, ALFA) |
| ***CNGB3:*c.1163-1205G>A** | rs1000861056 | T=0.000072 (19/264690, TOPMED)  **T=0.000021 (3/140068, gnomAD)**  T=0.00000 (0/14050, ALFA) |

**Supplementary Table 4:** Ratio (%) of genomic insertions upon editing to total reads obtained by high-throughput sequencing for the USH2A:.7595-2144A>G deep-intronic variants. Results refer to the sorted patient-derived fibroblasts. Results are presented as mean (%) ± SD of n=3 biological replicates.

|  | **EDCas9** | | **Cas9** | |
| --- | --- | --- | --- | --- |
| **Insertion size (bp)** | **1** | **2** | **1** | **2** |
| gRNA1 | 0.04±0.04 | 0.32±0.32 | 4.6±3.0 | 7.3±6.5 |
| gRNA3 | 0.17±0.15 | 0.002±0.003 | 4.7±3.2 | 0.51±0.47 |
| gRNA5 | 0.14±0.16 | 0.02±0.04 | 12.9±9.1 | 0.95±0.58 |
| gRNA6 | 0.05±0.05 | 0.003±0.005 | 6.5±3.0 | 0.85±1.04 |

**Supplementary Table 5:** List of primers and gRNA oligos used

| **Name** | **Sequence (5’ – 3’)** | **Use** |
| --- | --- | --- |
| **Primers** | | |
| Infusion-minigene-USH2A_F | ATGGGGTACGGGATCACCAGCGCACACACCCTTTCCAATATA | Primers to amplify and clone the *USH2A* fragment into pSPL3 |
| Infusion-minigene-USH2A_R | AGCGGCCGCTCGAGCTCCAGTGAGTGTGTGTATGCGATTCAG |  |
| Infusion-3xFlag-NLS_F | TCACTTTTTTTCAGGTTGGACCGGTGCC | Primers to amplify and clone 3xFLAG-NLS (EDCas9) |
| Infusion-3xFlag-NLS_R | TCGGCCCGGGGTGCCTCGGAGGCTGCTGGGACTCCGTGGATA |  |
| Infusion-TREX2-Linker/P2A_F | TCCGAGGCACCCCGGGCCGAGA | Primers to amplify and clone TREX2 (EDCas9 – EDCas9-P2A) |
| Infusion-TREX2-Linker_R | GCTGCCGCCTCCTCCGGCCTCCAGGCTGG |  |
| Infusion-TREX2-P2A_R | TCCTCGCCCTTGCTCACCGGTCCAGGATTCTCTTCGACATCTCCG |  |
| Infusion-SpCas9FR(P2A)_F | TCGAAGAGAATCCTGGACCG GACAAGAAGTACAGCATCGGCCTGG | Primers to amplify and clone part of SpCas9 (EDCas9 – EDCas9-P2A) |
| Infusion-SpCas9(Linker)FR_F | GGAGGAGGCGGCAGCGACAAGAAGTACAGCATCGGCCTGG |  |
| Infusion-SpCas9FR_R | AGTGTCAGGGTCAGCACGAT |  |
| USH2A_EX40_F | AATGGATTTGGCAGTGCACATA | Splicing assay in fibroblasts |
| USH2A_EX41_R | CCTTTTGAAGTGCAGGCTTCTA |  |
| pSPL3seqcDNA_F | TGGACAACCTCAAAGGCACC | Splicing assay by minigene in HEK293T for USH2A |
| pSPL3seqcDNA_R | AGTGAATTGGTCGAATGGATC |  |
| ABCA4_EX36_F | CTGCGTGATTTTCTCCATGTCC | Splicing assay by minigene in HEK293T for ABCA4 |
| ABCA4_EX37_R | GGTTTTCTGGAGAAGTGTAGGC |  |
| pSPL3_SA2_R | ATCTCAGTGGTATTTGTGAGC | pSPL3-specific primer for cDNA synthesis |
| USH2A-PE40_F | TGCAGTTGCAGGCCAGTTGATT | 1^st^ PCR amplification for NGS (gRNA1) |
| USH2A-PE40_R | TCTGTGATTGGGGGATAAGGCT |  |
| USH2A-PE40-2_F | CCTCTCCAGAATCACACAAG | 1^st^ PCR amplification for NGS (gRNA3, gRNA5, and gRNA6) |
| USH2A-PE40-2_R | CCTCTCTTCCCCAAAGAGAG |  |
| USH2A_PE40_NextAdpt_F | TCGTCGGCAGCGTCAGATGTGTATAAGAGACAGTGCAGTTGCAGGCCAGTTGATT | Primers to add Nextera Adapter (gRNA1) |
| USH2A_PE40_NextAdpt_R | GTCTCGTGGGCTCGGAGATGTGTATAAGAGACAGTCTGTGATTGGGGGATAAGGCT |  |
| USH2A_PE40_2-NextAdpt_F | TCGTCGGCAGCGTCAGATGTGTATAAGAGACAGCCTCTCCAGAATCACACAAG | Primers to add Nextera Adapter (gRNA3, gRNA5, and gRNA6) |
| USH2A_PE40_2-NextAdpt_R | GTCTCGTGGGCTCGGAGATGTGTATAAGAGACAGCCTCTCTTCCCCAAAGAGAG |  |
| CRISPR-insertcheck-rv2_R | CGCGCTAAAAACGGACTAGC | Primer to check gRNA cloning |
| pJET1.2_F | CGACTCACTATAGGGAGAGCGGC | Primers to sequence sub-cloned PCR amplicons |
| pJET1.2_R | AAGAACATCGATTTTCCATGGCAG |  |
| Off-target_gRNA3_1_F | GACCCACCCTGACTTTTACAAG | Primers to amplify genomic DNA for off-target sites |
| Off-target_gRNA3_1_R | AGGATTCGTTTGACTGAGGCTA |  |
| Off-target_gRNA3_2_F | TGCTTAGATTATGTCTCCCCTCC |  |
| Off-target_gRNA3_2_R | CGGAAAAATCCCATGGAACAGA |  |
| Off-target_gRNA3_3_F | CTGCACTGTAAGTTCCTCAAGG |  |
| Off-target_gRNA3_3_R | CCCTGACTCTGTTCCAAAAAGG |  |
| Off-target_gRNA6_1_F | TAGATTCAGGCCCCAATCAGAG |  |
| Off-target_gRNA6_1_R | GACATAAGCAATGCGAGAGGAA |  |
| Off-target_gRNA6_2_F | TTAACAACTCAGTTCTGCAGCC |  |
| Off-target_gRNA6_2_R | GCTAACAGCAGAGAACCTGTTC |  |
| Off-target_gRNA6_3_F | CATTGCACCAGATTCAGGTCTC |  |
| Off-target_gRNA6_3_R | CGAAAGCTTTTCACATTCCGGA |  |
| **gRNA oligos** | | |
| gRNA1_ABCA4-557_F | CACCGACAGGAGGCTGATCTGGTGC | Forward and reverse gRNA oligos used to clone the gRNAs into the backbone editing plasmids. Underlined the sequence of the cloning adapter. |
| gRNA1_ABCA4-557_R | AAACGCACCAGATCAGCCTCCTGTC |  |
| gRNA2_ABCA4-557_F | CACCGAGAAAGATATACTTACAGG |  |
| gRNA2_ABCA4-557_R | AAACCCTGTAAGTATATCTTTCTC |  |
| gRNA3_ABCA4-557_F | CACCGATACTTACAGGAGGCTGATC |  |
| gRNA3_ABCA4-557_R | AAACGATCAGCCTCCTGTAAGTATC |  |
| gRNA4_ABCA4-557_F | CACCGATGAGAAAGATATACTTAC |  |
| gRNA4_ABCA4-557_R | AAACGTAAGTATATCTTTCTCATC |  |
| gRNA5_ABCA4-557_F | CACCGATATCTTTCTCATCCCGTTG |  |
| gRNA5_ABCA4-557_R | AAACCAACGGGATGAGAAAGATATC |  |
| gRNA1_ABCA4cluster_F | CACCGCATCAACCCCAATTTATTCT |  |
| gRNA1_ABCA4cluster_R | AAACAGAATAAATTGGGGTTGATGC |  |
| gRNA2_ABCA4cluster_F | CACCGGATAAGAGCATCAACTGC |  |
| gRNA2_ABCA4cluster_R | AAACGCAGTTGATGCTCTTATCC |  |
| gRNA3wt_ABCA4_F | CACCGCTCTCTCTTCTGTCTACACG |  |
| gRNA3wt_ABCA4_R | AAACCGTGTAGACAGAAGAGAGAGC |  |
| gRNA3mut_ABCA4_F | CACCGCTCTCTCTTCTGTCTAGACG |  |
| gRNA3mut_ABCA4_R | AAACCGTCTAGACAGAAGAGAGAGC |  |
| gRNA1_USH2A_F | CACCGTAACTTGTGTGATTCTGGAG |  |
| gRNA1_USH2A_R | AAACCTCCAGAATCACACAAGTTAC |  |
| gRNA2_USH2A_F | CACCGAACACCTCTCCTTTCCCA |  |
| gRNA2_USH2A_R | AAACTGGGAAAGGAGAGGTGTTC |  |
| gRNA3_USH2A_F | CACCGTAAAGATGATCTCTTACCTT |  |
| gRNA3_USH2A_R | AAACAAGGTAAGAGATCATCTTTAC |  |
| gRNA4_USH2A_F | CACCGATGATCTCTTACCTTGGGAA |  |
| gRNA4_USH2A_R | AAACTTCCCAAGGTAAGAGATCATC |  |
| gRNA5_USH2A_F | CACCGCTCTTACCTTGGGAAAGGAG |  |
| gRNA5_USH2A_R | AAACCTCCTTTCCCAAGGTAAGAGC |  |
| gRNA6_USH2A_F | CACCGTTAAAGATGATCTCTTACCT |  |
| gRNA6_USH2A_R | AAACAGGTAAGAGATCATCTTTAAC |  |

**Supplementary Table 6:** Plasmid sequences

**pSPL3-USH2A(PE40)**

GGTGTGGAAAGTCCCCAGGCTCCCCAGCAGGCAGAAGTATGCAAAGCATGCATCTCAATTAGTCAGCAACCAGGTGTGGAAAGTCCCCAGGCTCCCCAGCAGGCAGAAGTATGCAAAGCATGCATCTCAATTAGTCAGCAACCATAGTCCCGCCCCTAACTCCGCCCATCCCGCCCCTAACTCCGCCCAGTTCCGCCCATTCTCCGCCCCATGGCTGACTAATTTTTTTTATTTATGCAGAGGCCGAGGCCGCCTCGGCCTCTGAGCTATTCCAGAAGTAGTGAGGAGGCTTTTTTGGAGGCCTAGGCTTTTGCAAAAAGCTTGGACTGTGTTTACTTGCAATCCCCCAAAACAGACAGAATGGTGCATCTGTCCAGTGAGGAGAAGTCTGCGGTCACTGCCCTGTGGGGCAAGGTGAATGTGGAAGAAGTTGGTGGTGAGGCCCTGGGCAGGCTGCTGGTTGTCTACCCATGGACCCAGAGGTTCTTCGAGTCCTTTGGGGACCTGTCCTCTGCAAATGCTGTTATGAACAATCCTAAGGTGAAGGCTCATGGCAAGAAGGTGCTGGCTGCCTTCAGTGAGGGTCTGAGTCACCTGGACAACCTCAAAGGCACCTTTGCTAAGCTGAGTGAACTGCACTGTGACAAGCTGCACGCTCTAGAGTCGACCCAGCAGTAAGTAATACATGTAATGCAACCTATACAAATAGCAATAGTAGCATTAGTAGTAGCAATAATAATAGCAATAGTTGTGTGGTCCATAGTAATCATAGAATATAGGAAAATATTAAGACAAAGAAAAATAGACAGGTTAATTGATAGACTAATAGAAAGAGCAGAAGACAGTGGCAATGAGAGTGAAGGAGAAATATCAGCACTTGTGGAGATGGGGGTGGAGATGGGGCACCATGCTCCTTGGGATGTTGATGATCTGTAGTGCTACAGAAAAATTGTGGGTCACAGTCTATTATGGGGTACGGGATCACCAGCGCACACACCCTTTCCAATATACACACCTGAATATAAAACTAAATAAGCAGATTCCTCTGTTAACAATTTTATTACTCTATTTTAGGCTGGGGCTGAACTTTTGAAGCTGATGAGCAAAATAGTTTCCCATAAAAGATAATTTCATTTGCCAAAATGAAACTACTTTCCGTTATTAATGTTGCAATGAAATGGTTTGAAATATGAGTCTGTGATCTGGACAGAAAGGATGCTCCTGTTCCCAGCTCCAAGAGGGAGTTCTCTAAACAATTACACCCACGGAGAGCGCTGTAAGGTTGTATTTTCATCACCGGCCAGCTGAGCTGTGTGCCTGTCATACTCCCAGTCTCCCCTCCAGTGACAGCCGCATTGACAACATGTTATACTCCACATTGATTTTACAGTTAAGGAGCTGATTGTGATATATAGAGATTTTAAGAATTCAGCTGAGAGGCTGACTTTTGCCTTTTATCATTCCATTAAAATTTTATAACCATCTTTTTCATGTCATTTCATCCATGAATTCTTACTTTAGAAAGTGATAGCTACTGCTGTGAGCATAACTTTTCTCATGTTCAGAGCTTTTTATTGTATTAGCAAACTACCCCAATTATAGTTATTACTCATGTTTTACATAATTGTGGTGGCCCTTTCAACCATGCAGTTGCAGGCCAGTTGATTTGTATATAGAATTAGATGATTCGGCTTATCATTTTAAAGCACTAAATTGAAAGAGTGCCAGGAGTCAGGTTTTAACACTTCCCTAGCCAAAGGAGCTAATTAAGCTGCTTTCAGCTTCCTCTCCAGAATCACACAAGTTAAAGGACCCTTCTGCAACAAGAGCAGCGAATCTACTCAGCCAGAGCAGGAAGCTAATAAAATGTATGCTGGCTTTTAAGGGGGAAACAAATCATGAAATTGAAATTGAACACCTCTCCTTTCCCAAG(**G/A**)TAAGAGATCATCTTTAAGAAAAGGCTGTGTATTGTGGGGGTTTGAAGTGCAAGTTCATCTCATTATCATGGATGTTTCACCCATAATACTATCATCATATGCAGGAGAAATAAAAGCCTTATCCCCCAATCACAGAGAACAAACTGCATCATTGTTGTTTCTTTTCCCTGAAGATGCTCACAGTTTGATAGTTCACATATGAGGAATAATTATGCAAGACCCAGATGAAAACCCTAATAAAAAATATTCTGATTTTTAAGTGTATATTACTCTCTTTGGGGAAGAGAGGGTGAGATAAGGTTAGACATTAAAGACTTGAATCTCCACCATTGATTTTACGAAACCCAATCTCAGGGGATTTGGGGGTTTTTTTAGATTTCAAAACAACATTGATATCTTACAAGACTCGTACTTGCTTTTCTCTGGGAAATAGCCAAATCTAGATGACAAGCATAAATAAGAAAGGAGCTGTATCCCCATTTTGATATAATAATTGCTACTGCTACTGTATACCAGAGAGCAAAAGTAAGAAATGGCATAGTGGTTTTAAGAAAGCTTTAAAATGAAAGCTAACAAGAGACTGAAACGGAAAGTGTGATTTATACCATGACTGCTGGTTTTAGGGTTTTTTTTAGGGATTTCTCTTCAAATAGGATACATTACCAAATTGTCAGAAATAGCTTGATTTCAGAGTTAGTGCACTATAACATCTTTTGCTTCGAAAAGTGATTATGATTCTACCATCAGTCCAGTTGGTGACAAATACGTGGATCCTGTTGTGTGAAAGTGTACTCATATAACCAACTATTATTGCAGTCACTGGCCCTGCAAAGCAGTGGTTGCATTTAAAATTTCAACCTTCTTTGTAACACTGAATCGCATACACACACTCACTGGAGCTCGAGCGGCCGCTGCAGGATCCCAGATATCTGGTGATCCCGTACCTGTGTGGAAGGAAGCAACCACCACTCTATTTTGTGCATCAGATGCTAAAGCATATGATACAGAGGTACATAATGTTTGGGCCACACATGCCGGTGTACCCACAGACCCCAACCCACAAGAAGTAGTATTGGTAAATGTGACAGAAAATTTTAACATGTGGAAAAATGACATGGTAGAACAGATGCATGAGGATATAATCAGTTTATGGGATCAAAGCCTAAAGCCATGTGTAAAATTAACCCCACTCTGTGTTAGTTTAAAGTGCACTGATTTGAAGAATGATACTAATACCAATAGTAGTAGCGGGAGAATGATAATGGAGAAAGGAGAGATAAAAAACTGCTCTTTCAATATCAGCACAAGCATAAGAGGTAAGGTGCAGAAAGAATATGCATTTTTTTATAAACTTGATATAATACCAATAGATAATGATACTACCAGCTATACGTTGACAAGTTGTAACACCTCAGTCATTACACAGGCCTGTCCAAAGGTATCCTTTGAGCCAATTCCCATACATTATTGTGCCCCGGCTGGTTTTGCGATTCTAAAATGTAATAATAAGACGTTCAATGGAACAGGACCATGTACAAATGTCAGCACAGTACAATGTACACATGGAATTAGGCCAGTAGTATCAACTCAACTGCTGTTAAATGGCAGTCTAGCAGAAGAAGAGGTAGTAATTAGATCTGTCAATTTCACGGACAATGCTAAAACCATAATAGTACAGCTGAACACATCTGTAGAAATTAATTGTACAAGACCCAACAACAATACAAGAAAAAAAATCCGTATCCAGAGGGGACCAGGGAGAGCATTTGTTACAATAGGAAAAATAGGAAATATGAGACAAGCACATTGTAACATTAGTAGAGCAAAATGGAATGCCACTTTAAAACAGATAGCTAGCAAATTAAGAGAACAATTTGGAAATAATAAAACAATAATCTTTAAGCAATCCTCAGGAGGGGACCCAGAAATTGTAACGCACAGTTTTAATTGTGGAGGGGAATTTTTCTACTGTAATTCAACACAACTGTTTAATAGTACTTGGTTTAATAGTACTTGGAGTACTGAAGGGTCAAATAACACTGAAGGAAGTGACACAATCACACTCCCATGCAGAATAAAACAATTTATAAACATGTGGCAGGAAGTAGGAAAAGCAATGTATGCCCCTCCCATCAGCGGACAAATTAGATGTTCATCAAATATTACAGGGCTGCTATTAACAAGAGATGGTGGTAATAACAACAATGGGTCCGAGATCTTCAGACCTGGAGGAGGAGATATGAGGGACAATTGGAGAAGTGAATTATATAAATATAAAGTAGTAAAAATTGAACCATTAGGAGTAGCACCCACCAAGGCAAAGAGAAGAGTGGTGCAGAGAGAAAAAAGAGCAGTGGGAATAGGAGCTTTGTTCCTTGGGTTCTTGGGAGCAGCAGGAAGCACTATGGGCGCAGCGTCAATGACGCTGACGGTACAGGCCAGACAATTATTGTCTGGTATAGTGCAGCAGCAGAACAATTTGCTGAGGGCTATTGAGGCGCAACAGCATCTGTTGCAACTCACAGTCTGGGGCATCAAGCAGCTCCAGGCAAGAATCCTGGCTGTGGAAAGATACCTAAAGGATCAACAGCTCCTGGGGATTTGGGGTTGCTCTGGAAAACTACTTTGCACCACTGCTGTGCCTTGGAATGCTAGTTGGAGTAATAAATCTCTGGAACAGATTTGGAATCACACGACGTGGATGGAGTGGGACAGAGAAATTAACAATTACACAAGCTTAATACACTCCTTAATTGAAGAATCGCAAAACCAGCAAGAAAAGAATGAACAAGAATTATTGGAATTAGATAAATGGGCAAGTTTGTGGAATTGGTTTAACATAACAAATTGGCTGTGGTATATAAAATTATTCATAATGATAGTAGGAGGCTTGGTAGGTTTAAGAATAGTTTTTGCTGTACTTTCTGTAGTGAATAGAGTTAGGCAGGGATATTCACCATTATCGTTTCAGACCTGGAGATCTCCCGAGGGGACCCGACAGGCCCGAAGGAATAGAAGAAGAAGGTGGAGAGAGAGACAGAGACAGATCCATTCGACCAATTCACTCCTCAGGTGCAGGCTGCCTATCAGAAGGTGGTGGCTGGTGTGGCCAATGCCCTGGCTCACAAATACCACTGAGATCCAGACATGATAAGATACATTGATGAGTTTGGACAAACCACAACTAGAATGCAGTGAAAAAAATGCTTTATTTGTGAAATTTGTGATGCTATTGCTTTATTTGTAACCATTATAAGCTGCAATAAACAAGTTAACAACAACAATTGCATTCATTTTATGTTTCAGGTTCAGGGGGAGGTGTGGGAGGTTTTTTAAAGCAAGTAAAACCTCTACAAATGTGGTATGGCTGATTATGATCCCCAGGAAGCTCCTCTGTGTCCTCATAAACCCTAACCTCCTCTACTTGAGAGGACATTCCAATCATAGGCTGCCCATCCACCCTCTGTGTCCTCCTGTTAATTAGGTCACTTAACAAAAAGGAAATTGGGTAGGGGTTTTTCACAGACCGCTTTCTAAGGGTAATTTTAAAATATCTGGGAAGTCCCTTCCACTGCTGTGTTCCAGAAGTGTTGGTAAACAGCCCACAAATGTCAACAGCAGAAACATACAAGCTGTCAGCTTTGCACAAGGGCCCAACACCCTGCTCATCAAGAAGCACTGTGGTTGCTGTGTTAGTAATGTGCAAAACAGGAGGCACATTTTCCCCACCTGTGTAGGTTCCAAAATATCTAGTGTTTTCATTTTTACTTGGATCAGGAACCCAGCACTCCACTGGATAAGCATTATCCTTATCCAAAACAGCCTTGTGGTCAGTGTTCATCTGCTGACTGTCAACTGTAGCATTTTTTGGGGTTACAGTTTGAGCAGGATATTTGGTCCTGTAGTTTGCTAACACACCCCAGGTGGCACTTTTCGGGGAAATGTGCGCGGAACCCCTATTTGTTTATTTTTCTAAATACATTCAAATATGTATCCGCTCATGAGACAATAACCCTGATAAATGCTTCAATAATATTGAAAAAGGAAGAGTATGAGTATTCAACATTTCCGTGTCGCCCTTATTCCCTTTTTTGCGGCATTTTGCCTTCCTGTTTTTGCTCACCCAGAAACGCTGGTGAAAGTAAAAGATGCTGAAGATCAGTTGGGTGCACGAGTGGGTTACATCGAACTGGATCTCAACAGCGGTAAGATCCTTGAGAGTTTTCGCCCCGAAGAACGTTTTCCAATGATGAGCACTTTTAAAGTTCTGCTATGTGGCGCGGTATTATCCCGTATTGACGCCGGGCAAGAGCAACTCGGTCGCCGCATACACTATTCTCAGAATGACTTGGTTGAGTACTCACCAGTCACAGAAAAGCATCTTACGGATGGCATGACAGTAAGAGAATTATGCAGTGCTGCCATAACCATGAGTGATAACACTGCGGCCAACTTACTTCTGACAACGATCGGAGGACCGAAGGAGCTAACCGCTTTTTTGCACAACATGGGGGATCATGTAACTCGCCTTGATCGTTGGGAACCGGAGCTGAATGAAGCCATACCAAACGACGAGCGTGACACCACGATGCCTGTAGCAATGGCAACAACGTTGCGCAAACTATTAACTGGCGAACTACTTACTCTAGCTTCCCGGCAACAATTAATAGACTGGATGGAGGCGGATAAAGTTGCAGGACCACTTCTGCGCTCGGCCCTTCCGGCTGGCTGGTTTATTGCTGATAAATCTGGAGCCGGTGAGCGTGGGTCTCGCGGTATCATTGCAGCACTGGGGCCAGATGGTAAGCCCTCCCGTATCGTAGTTATCTACACGACGGGGAGTCAGGCAACTATGGATGAACGAAATAGACAGATCGCTGAGATAGGTGCCTCACTGATTAAGCATTGGTAACTGTCAGACCAAGTTTACTCATATATACTTTAGATTGATTTAAAACTTCATTTTTAATTTAAAAGGATCTAGGTGAAGATCCTTTTTGATAATCTCATGACCAAAATCCTTAACGGTGAGTTTTCGTTCCACTGAGCGTCAGACCCCGTAGAAAAGATCAAAGGATCTTCTTGAGATCCTTTTTTTCTGCGCGTAATCTGCTGCTTGCAAACAAAAAAACCACCGCTACCAGCGGTGGTTTGTTTGCCGGATCAAGAGCTACCAACTCTTTTTCCGAAGGTAACTGGCTTCAGCAGAGCGCAGATACCAAATACTGTCCTTCTAGTGTAGCCGTAGTTAGGCCACCACTTCAAGAACTCTGTAGCACCGCCTACATACCTCGCTCTGCTAATCCTGTTACCAGTGGCTGCTGCCAGTGGCGATAAGTCGTGTCTTACCGGGTTGGACTCAAGACGATAGTTACCGGATAAGGCGCAGCGGTCGGGCTGAACGGGGGGTTCGTGCACACAGCCCAGCTTGGAGCGAACGACCTACACCGAACTGAGATACCTACAGCGCGAGCATTGAGAAAGCGCCACGCTTCCCGAAGGGAGAAAGGCGGACAGGTATCCGGTAAGCGGCAGGGTCGGAACAGGAGAGCGCACGAGGGAGCTTCCAGGGGGAAACGCCTGGTATCTTTATAGTCCTGTCGGGTTTCGCCACCTCTGACTTGAGCGTCGATTTTTGTGATGCTCGTCAGGGGGGCGGAGCCTATGGAAAAACGCCAGCAACGCGGCCTTTTTACGGTTCCTGGCCTTTTGCTGGCCTTTTGCTCACATGTTCTTTCCTGCGTTATCCCCTGATTCTGTGGATAACCGTATTACCGCCTTTGAGTGAGCTGATACCGCTCGCCGCAGCCGAACGACCGAGCGCAGCGAGTCAGTGAGCGAGGAAGCGGAAGAGCGCCCAATACGCAAACCGCCTCTCCCCGCGCGTTGGCCGATTCATTAATGCAGCTGTGGAATGTGTGTCAGTTAG

Legend: SV40 Promoter, TAT1-HIV, USH2A-partial intron40, PE40 (152 bp), **USH2A:c.7595-2144A/G**,

**EDCas9** (U6-gRNA_CAG-NLS-TREX2-Linker-SpCas9-NLS-T2A-EGFP

GAGGGCCTATTTCCCATGATTCCTTCATATTTGCATATACGATACAAGGCTGTTAGAGAGATAATTGGAATTAATTTGACTGTAAACACAAAGATATTAGTACAAAATACGTGACGTAGAAAGTAATAATTTCTTGGGTAGTTTGCAGTTTTAAAATTATGTTTTAAAATGGACTATCATATGCTTACCGTAACTTGAAAGTATTTCGATTTCTTGGCTTTATATATCTTGTGGAAAGGACGAAACACCGGGTCTTCGAGAAGACACGTTTTAGAGCTAGAAATAGCAAGTTAAAATAAGGCTAGTCCGTTATCAACTTGAAAAAGTGGCACCGAGTCGGTGCTTTTTTGTTTTAGAGCTAGAAATAGCAAGTTAAAATAAGGCTAGTCCGTTTTTAGCGCGTGCGCCAATTCTGCAGACAAATGGCTCTAGAGGTACCCGTTACATAACTTACGGTAAATGGCCCGCCTGGCTGACCGCCCAACGACCCCCGCCCATTGACGTCAATAGTAACGCCAATAGGGACTTTCCATTGACGTCAATGGGTGGAGTATTTACGGTAAACTGCCCACTTGGCAGTACATCAAGTGTATCATATGCCAAGTACGCCCCCTATTGACGTCAATGACGGTAAATGGCCCGCCTGGCATTGTGCCCAGTACATGACCTTATGGGACTTTCCTACTTGGCAGTACATCTACGTATTAGTCATCGCTATTACCATGGTCGAGGTGAGCCCCACGTTCTGCTTCACTCTCCCCATCTCCCCCCCCTCCCCACCCCCAATTTTGTATTTATTTATTTTTTAATTATTTTGTGCAGCGATGGGGGCGGGGGGGGGGGGGGGGCGCGCGCCAGGCGGGGCGGGGCGGGGCGAGGGGCGGGGCGGGGCGAGGCGGAGAGGTGCGGCGGCAGCCAATCAGAGCGGCGCGCTCCGAAAGTTTCCTTTTATGGCGAGGCGGCGGCGGCGGCGGCCCTATAAAAAGCGAAGCGCGCGGCGGGCGGGAGTCGCTGCGCGCTGCCTTCGCCCCGTGCCCCGCTCCGCCGCCGCCTCGCGCCGCCCGCCCCGGCTCTGACTGACCGCGTTACTCCCACAGGTGAGCGGGCGGGACGGCCCTTCTCCTCCGGGCTGTAATTAGCTGAGCAAGAGGTAAGGGTTTAAGGGATGGTTGGTTGGTGGGGTATTAATGTTTAATTACCTGGAGCACCTGCCTGAAATCACTTTTTTTCAGGTTGGACCGGTGCCACCATGGACTATAAGGACCACGACGGAGACTACAAGGATCATGATATTGATTACAAAGACGATGACGATAAGATGGCCCCAAAGAAGAAGCGGAAGGTCGGTATCCACGGAGTCCCAGCAGCCTCCGAGGCACCCCGGGCCGAGACCTTTGTGTTCCTGGACCTGGAAGCCACTGGGCTCCCCAGTGTGGAGCCCGAGATTGCCGAGCTGTCCCTCTTTGCTGTCCACCGCTCCTCCCTGGAGAACCCGGAGCACGACGAGTCTGGTGCCCTAGTATTGCCCCGGGTCCTGGACAAGCTCACGCTGTGCATGTGCCCGGAGCGCCCCTTCACTGCCAAGGCCAGCGAGATCACCGGCCTGAGCAGTGAGGGCCTGGCGCGATGCCGGAAGGCTGGCTTTGATGGCGCCGTGGTGCGGACGCTGCAGGCCTTCCTGAGCCGCCAGGCAGGGCCCATCTGCCTTGTGGCCCACAATGGCTTTGATTATGATTTCCCCCTGCTGTGTGCCGAGCTGCGGCGCCTGGGTGCTCGCCTGCCTCGGGACACTGTCTGCCTGGACACGCTGCCAGCCCTGCGGGGCCTGGACCGCGCCCACAGCCACGGCACACGAGCTAGAGGCAGACAGGGTTACAGCCTCGGCAGCCTCTTCCACCGCTACTTCCGGGCAGAGCCAAGCGCAGCCCACTCAGCCGAGGGCGACGTGCACACCCTGCTCCTGATCTTCCTGCACCGCGCCGCAGAGCTGCTCGCCTGGGCCGATGAGCAGGCCCGTGGGTGGGCCCACATCGAGCCCATGTACTTGCCGCCTGATGACCCCAGCCTGGAGGCCGGAGGAGGCGGCAGCGACAAGAAGTACAGCATCGGCCTGGACATCGGCACCAACTCTGTGGGCTGGGCCGTGATCACCGACGAGTACAAGGTGCCCAGCAAGAAATTCAAGGTGCTGGGCAACACCGACCGGCACAGCATCAAGAAGAACCTGATCGGAGCCCTGCTGTTCGACAGCGGCGAAACAGCCGAGGCCACCCGGCTGAAGAGAACCGCCAGAAGAAGATACACCAGACGGAAGAACCGGATCTGCTATCTGCAAGAGATCTTCAGCAACGAGATGGCCAAGGTGGACGACAGCTTCTTCCACAGACTGGAAGAGTCCTTCCTGGTGGAAGAGGATAAGAAGCACGAGCGGCACCCCATCTTCGGCAACATCGTGGACGAGGTGGCCTACCACGAGAAGTACCCCACCATCTACCACCTGAGAAAGAAACTGGTGGACAGCACCGACAAGGCCGACCTGCGGCTGATCTATCTGGCCCTGGCCCACATGATCAAGTTCCGGGGCCACTTCCTGATCGAGGGCGACCTGAACCCCGACAACAGCGACGTGGACAAGCTGTTCATCCAGCTGGTGCAGACCTACAACCAGCTGTTCGAGGAAAACCCCATCAACGCCAGCGGCGTGGACGCCAAGGCCATCCTGTCTGCCAGACTGAGCAAGAGCAGACGGCTGGAAAATCTGATCGCCCAGCTGCCCGGCGAGAAGAAGAATGGCCTGTTCGGAAACCTGATTGCCCTGAGCCTGGGCCTGACCCCCAACTTCAAGAGCAACTTCGACCTGGCCGAGGATGCCAAACTGCAGCTGAGCAAGGACACCTACGACGACGACCTGGACAACCTGCTGGCCCAGATCGGCGACCAGTACGCCGACCTGTTTCTGGCCGCCAAGAACCTGTCCGACGCCATCCTGCTGAGCGACATCCTGAGAGTGAACACCGAGATCACCAAGGCCCCCCTGAGCGCCTCTATGATCAAGAGATACGACGAGCACCACCAGGACCTGACCCTGCTGAAAGCTCTCGTGCGGCAGCAGCTGCCTGAGAAGTACAAAGAGATTTTCTTCGACCAGAGCAAGAACGGCTACGCCGGCTACATTGACGGCGGAGCCAGCCAGGAAGAGTTCTACAAGTTCATCAAGCCCATCCTGGAAAAGATGGACGGCACCGAGGAACTGCTCGTGAAGCTGAACAGAGAGGACCTGCTGCGGAAGCAGCGGACCTTCGACAACGGCAGCATCCCCCACCAGATCCACCTGGGAGAGCTGCACGCCATTCTGCGGCGGCAGGAAGATTTTTACCCATTCCTGAAGGACAACCGGGAAAAGATCGAGAAGATCCTGACCTTCCGCATCCCCTACTACGTGGGCCCTCTGGCCAGGGGAAACAGCAGATTCGCCTGGATGACCAGAAAGAGCGAGGAAACCATCACCCCCTGGAACTTCGAGGAAGTGGTGGACAAGGGCGCTTCCGCCCAGAGCTTCATCGAGCGGATGACCAACTTCGATAAGAACCTGCCCAACGAGAAGGTGCTGCCCAAGCACAGCCTGCTGTACGAGTACTTCACCGTGTATAACGAGCTGACCAAAGTGAAATACGTGACCGAGGGAATGAGAAAGCCCGCCTTCCTGAGCGGCGAGCAGAAAAAGGCCATCGTGGACCTGCTGTTCAAGACCAACCGGAAAGTGACCGTGAAGCAGCTGAAAGAGGACTACTTCAAGAAAATCGAGTGCTTCGACTCCGTGGAAATCTCCGGCGTGGAAGATCGGTTCAACGCCTCCCTGGGCACATACCACGATCTGCTGAAAATTATCAAGGACAAGGACTTCCTGGACAATGAGGAAAACGAGGACATTCTGGAAGATATCGTGCTGACCCTGACACTGTTTGAGGACAGAGAGATGATCGAGGAACGGCTGAAAACCTATGCCCACCTGTTCGACGACAAAGTGATGAAGCAGCTGAAGCGGCGGAGATACACCGGCTGGGGCAGGCTGAGCCGGAAGCTGATCAACGGCATCCGGGACAAGCAGTCCGGCAAGACAATCCTGGATTTCCTGAAGTCCGACGGCTTCGCCAACAGAAACTTCATGCAGCTGATCCACGACGACAGCCTGACCTTTAAAGAGGACATCCAGAAAGCCCAGGTGTCCGGCCAGGGCGATAGCCTGCACGAGCACATTGCCAATCTGGCCGGCAGCCCCGCCATTAAGAAGGGCATCCTGCAGACAGTGAAGGTGGTGGACGAGCTCGTGAAAGTGATGGGCCGGCACAAGCCCGAGAACATCGTGATCGAAATGGCCAGAGAGAACCAGACCACCCAGAAGGGACAGAAGAACAGCCGCGAGAGAATGAAGCGGATCGAAGAGGGCATCAAAGAGCTGGGCAGCCAGATCCTGAAAGAACACCCCGTGGAAAACACCCAGCTGCAGAACGAGAAGCTGTACCTGTACTACCTGCAGAATGGGCGGGATATGTACGTGGACCAGGAACTGGACATCAACCGGCTGTCCGACTACGATGTGGACCATATCGTGCCTCAGAGCTTTCTGAAGGACGACTCCATCGACAACAAGGTGCTGACCAGAAGCGACAAGAACCGGGGCAAGAGCGACAACGTGCCCTCCGAAGAGGTCGTGAAGAAGATGAAGAACTACTGGCGGCAGCTGCTGAACGCCAAGCTGATTACCCAGAGAAAGTTCGACAATCTGACCAAGGCCGAGAGAGGCGGCCTGAGCGAACTGGATAAGGCCGGCTTCATCAAGAGACAGCTGGTGGAAACCCGGCAGATCACAAAGCACGTGGCACAGATCCTGGACTCCCGGATGAACACTAAGTACGACGAGAATGACAAGCTGATCCGGGAAGTGAAAGTGATCACCCTGAAGTCCAAGCTGGTGTCCGATTTCCGGAAGGATTTCCAGTTTTACAAAGTGCGCGAGATCAACAACTACCACCACGCCCACGACGCCTACCTGAACGCCGTCGTGGGAACCGCCCTGATCAAAAAGTACCCTAAGCTGGAAAGCGAGTTCGTGTACGGCGACTACAAGGTGTACGACGTGCGGAAGATGATCGCCAAGAGCGAGCAGGAAATCGGCAAGGCTACCGCCAAGTACTTCTTCTACAGCAACATCATGAACTTTTTCAAGACCGAGATTACCCTGGCCAACGGCGAGATCCGGAAGCGGCCTCTGATCGAGACAAACGGCGAAACCGGGGAGATCGTGTGGGATAAGGGCCGGGATTTTGCCACCGTGCGGAAAGTGCTGAGCATGCCCCAAGTGAATATCGTGAAAAAGACCGAGGTGCAGACAGGCGGCTTCAGCAAAGAGTCTATCCTGCCCAAGAGGAACAGCGATAAGCTGATCGCCAGAAAGAAGGACTGGGACCCTAAGAAGTACGGCGGCTTCGACAGCCCCACCGTGGCCTATTCTGTGCTGGTGGTGGCCAAAGTGGAAAAGGGCAAGTCCAAGAAACTGAAGAGTGTGAAAGAGCTGCTGGGGATCACCATCATGGAAAGAAGCAGCTTCGAGAAGAATCCCATCGACTTTCTGGAAGCCAAGGGCTACAAAGAAGTGAAAAAGGACCTGATCATCAAGCTGCCTAAGTACTCCCTGTTCGAGCTGGAAAACGGCCGGAAGAGAATGCTGGCCTCTGCCGGCGAACTGCAGAAGGGAAACGAACTGGCCCTGCCCTCCAAATATGTGAACTTCCTGTACCTGGCCAGCCACTATGAGAAGCTGAAGGGCTCCCCCGAGGATAATGAGCAGAAACAGCTGTTTGTGGAACAGCACAAGCACTACCTGGACGAGATCATCGAGCAGATCAGCGAGTTCTCCAAGAGAGTGATCCTGGCCGACGCTAATCTGGACAAAGTGCTGTCCGCCTACAACAAGCACCGGGATAAGCCCATCAGAGAGCAGGCCGAGAATATCATCCACCTGTTTACCCTGACCAATCTGGGAGCCCCTGCCGCCTTCAAGTACTTTGACACCACCATCGACCGGAAGAGGTACACCAGCACCAAAGAGGTGCTGGACGCCACCCTGATCCACCAGAGCATCACCGGCCTGTACGAGACACGGATCGACCTGTCTCAGCTGGGAGGCGACAAAAGGCCGGCGGCCACGAAAAAGGCCGGCCAGGCAAAAAAGAAAAAGGAATTCGGCAGTGGAGAGGGCAGAGGAAGTCTGCTAACATGCGGTGACGTCGAGGAGAATCCTGGCCCAGTGAGCAAGGGCGAGGAGCTGTTCACCGGGGTGGTGCCCATCCTGGTCGAGCTGGACGGCGACGTAAACGGCCACAAGTTCAGCGTGTCCGGCGAGGGCGAGGGCGATGCCACCTACGGCAAGCTGACCCTGAAGTTCATCTGCACCACCGGCAAGCTGCCCGTGCCCTGGCCCACCCTCGTGACCACCCTGACCTACGGCGTGCAGTGCTTCAGCCGCTACCCCGACCACATGAAGCAGCACGACTTCTTCAAGTCCGCCATGCCCGAAGGCTACGTCCAGGAGCGCACCATCTTCTTCAAGGACGACGGCAACTACAAGACCCGCGCCGAGGTGAAGTTCGAGGGCGACACCCTGGTGAACCGCATCGAGCTGAAGGGCATCGACTTCAAGGAGGACGGCAACATCCTGGGGCACAAGCTGGAGTACAACTACAACAGCCACAACGTCTATATCATGGCCGACAAGCAGAAGAACGGCATCAAGGTGAACTTCAAGATCCGCCACAACATCGAGGACGGCAGCGTGCAGCTCGCCGACCACTACCAGCAGAACACCCCCATCGGCGACGGCCCCGTGCTGCTGCCCGACAACCACTACCTGAGCACCCAGTCCGCCCTGAGCAAAGACCCCAACGAGAAGCGCGATCACATGGTCCTGCTGGAGTTCGTGACCGCCGCCGGGATCACTCTCGGCATGGACGAGCTGTACAAGGAATTCTAACTAGAGCTCGCTGATCAGCCTCGACTGTGCCTTCTAGTTGCCAGCCATCTGTTGTTTGCCCCTCCCCCGTGCCTTCCTTGACCCTGGAAGGTGCCACTCCCACTGTCCTTTCCTAATAAAATGAGGAAATTGCATCGCATTGTCTGAGTAGGTGTCATTCTATTCTGGGGGGTGGGGTGGGGCAGGACAGCAAGGGGGAGGATTGGGAAGAGAATAGCAGGCATGCTGGGGAGCGGCCGCAGGAACCCCTAGTGATGGAGTTGGCCACTCCCTCTCTGCGCGCTCGCTCGCTCACTGAGGCCGGGCGACCAAAGGTCGCCCGACGCCCGGGCTTTGCCCGGGCGGCCTCAGTGAGCGAGCGAGCGCGCAGCTGCCTGCAGGGGCGCCTGATGCGGTATTTTCTCCTTACGCATCTGTGCGGTATTTCACACCGCATACGTCAAAGCAACCATAGTACGCGCCCTGTAGCGGCGCATTAAGCGCGGCGGGTGTGGTGGTTACGCGCAGCGTGACCGCTACACTTGCCAGCGCCTTAGCGCCCGCTCCTTTCGCTTTCTTCCCTTCCTTTCTCGCCACGTTCGCCGGCTTTCCCCGTCAAGCTCTAAATCGGGGGCTCCCTTTAGGGTTCCGATTTAGTGCTTTACGGCACCTCGACCCCAAAAAACTTGATTTGGGTGATGGTTCACGTAGTGGGCCATCGCCCTGATAGACGGTTTTTCGCCCTTTGACGTTGGAGTCCACGTTCTTTAATAGTGGACTCTTGTTCCAAACTGGAACAACACTCAACTCTATCTCGGGCTATTCTTTTGATTTATAAGGGATTTTGCCGATTTCGGTCTATTGGTTAAAAAATGAGCTGATTTAACAAAAATTTAACGCGAATTTTAACAAAATATTAACGTTTACAATTTTATGGTGCACTCTCAGTACAATCTGCTCTGATGCCGCATAGTTAAGCCAGCCCCGACACCCGCCAACACCCGCTGACGCGCCCTGACGGGCTTGTCTGCTCCCGGCATCCGCTTACAGACAAGCTGTGACCGTCTCCGGGAGCTGCATGTGTCAGAGGTTTTCACCGTCATCACCGAAACGCGCGAGACGAAAGGGCCTCGTGATACGCCTATTTTTATAGGTTAATGTCATGATAATAATGGTTTCTTAGACGTCAGGTGGCACTTTTCGGGGAAATGTGCGCGGAACCCCTATTTGTTTATTTTTCTAAATACATTCAAATATGTATCCGCTCATGAGACAATAACCCTGATAAATGCTTCAATAATATTGAAAAAGGAAGAGTATGAGTATTCAACATTTCCGTGTCGCCCTTATTCCCTTTTTTGCGGCATTTTGCCTTCCTGTTTTTGCTCACCCAGAAACGCTGGTGAAAGTAAAAGATGCTGAAGATCAGTTGGGTGCACGAGTGGGTTACATCGAACTGGATCTCAACAGCGGTAAGATCCTTGAGAGTTTTCGCCCCGAAGAACGTTTTCCAATGATGAGCACTTTTAAAGTTCTGCTATGTGGCGCGGTATTATCCCGTATTGACGCCGGGCAAGAGCAACTCGGTCGCCGCATACACTATTCTCAGAATGACTTGGTTGAGTACTCACCAGTCACAGAAAAGCATCTTACGGATGGCATGACAGTAAGAGAATTATGCAGTGCTGCCATAACCATGAGTGATAACACTGCGGCCAACTTACTTCTGACAACGATCGGAGGACCGAAGGAGCTAACCGCTTTTTTGCACAACATGGGGGATCATGTAACTCGCCTTGATCGTTGGGAACCGGAGCTGAATGAAGCCATACCAAACGACGAGCGTGACACCACGATGCCTGTAGCAATGGCAACAACGTTGCGCAAACTATTAACTGGCGAACTACTTACTCTAGCTTCCCGGCAACAATTAATAGACTGGATGGAGGCGGATAAAGTTGCAGGACCACTTCTGCGCTCGGCCCTTCCGGCTGGCTGGTTTATTGCTGATAAATCTGGAGCCGGTGAGCGTGGAAGCCGCGGTATCATTGCAGCACTGGGGCCAGATGGTAAGCCCTCCCGTATCGTAGTTATCTACACGACGGGGAGTCAGGCAACTATGGATGAACGAAATAGACAGATCGCTGAGATAGGTGCCTCACTGATTAAGCATTGGTAACTGTCAGACCAAGTTTACTCATATATACTTTAGATTGATTTAAAACTTCATTTTTAATTTAAAAGGATCTAGGTGAAGATCCTTTTTGATAATCTCATGACCAAAATCCCTTAACGTGAGTTTTCGTTCCACTGAGCGTCAGACCCCGTAGAAAAGATCAAAGGATCTTCTTGAGATCCTTTTTTTCTGCGCGTAATCTGCTGCTTGCAAACAAAAAAACCACCGCTACCAGCGGTGGTTTGTTTGCCGGATCAAGAGCTACCAACTCTTTTTCCGAAGGTAACTGGCTTCAGCAGAGCGCAGATACCAAATACTGTTCTTCTAGTGTAGCCGTAGTTAGGCCACCACTTCAAGAACTCTGTAGCACCGCCTACATACCTCGCTCTGCTAATCCTGTTACCAGTGGCTGCTGCCAGTGGCGATAAGTCGTGTCTTACCGGGTTGGACTCAAGACGATAGTTACCGGATAAGGCGCAGCGGTCGGGCTGAACGGGGGGTTCGTGCACACAGCCCAGCTTGGAGCGAACGACCTACACCGAACTGAGATACCTACAGCGTGAGCTATGAGAAAGCGCCACGCTTCCCGAAGGGAGAAAGGCGGACAGGTATCCGGTAAGCGGCAGGGTCGGAACAGGAGAGCGCACGAGGGAGCTTCCAGGGGGAAACGCCTGGTATCTTTATAGTCCTGTCGGGTTTCGCCACCTCTGACTTGAGCGTCGATTTTTGTGATGCTCGTCAGGGGGGCGGAGCCTATGGAAAAACGCCAGCAACGCGGCCTTTTTACGGTTCCTGGCCTTTTGCTGGCCTTTTGCTCACATGT

Legend: U6 Promoter, gRNA scaffold, CAG promoter, TREX2, Linker, SpCas9, NLS signals, T2A signal, EGFP

**EDCas9-P2A** (U6-gRNA_CAG-NLS-SpCas9-P2A-TREX2-T2A-EGFP

GAGGGCCTATTTCCCATGATTCCTTCATATTTGCATATACGATACAAGGCTGTTAGAGAGATAATTGGAATTAATTTGACTGTAAACACAAAGATATTAGTACAAAATACGTGACGTAGAAAGTAATAATTTCTTGGGTAGTTTGCAGTTTTAAAATTATGTTTTAAAATGGACTATCATATGCTTACCGTAACTTGAAAGTATTTCGATTTCTTGGCTTTATATATCTTGTGGAAAGGACGAAACACCGGGTCTTCGAGAAGACACGTTTTAGAGCTAGAAATAGCAAGTTAAAATAAGGCTAGTCCGTTATCAACTTGAAAAAGTGGCACCGAGTCGGTGCTTTTTTGTTTTAGAGCTAGAAATAGCAAGTTAAAATAAGGCTAGTCCGTTTTTAGCGCGTGCGCCAATTCTGCAGACAAATGGCTCTAGAGGTACCCGTTACATAACTTACGGTAAATGGCCCGCCTGGCTGACCGCCCAACGACCCCCGCCCATTGACGTCAATAGTAACGCCAATAGGGACTTTCCATTGACGTCAATGGGTGGAGTATTTACGGTAAACTGCCCACTTGGCAGTACATCAAGTGTATCATATGCCAAGTACGCCCCCTATTGACGTCAATGACGGTAAATGGCCCGCCTGGCATTGTGCCCAGTACATGACCTTATGGGACTTTCCTACTTGGCAGTACATCTACGTATTAGTCATCGCTATTACCATGGTCGAGGTGAGCCCCACGTTCTGCTTCACTCTCCCCATCTCCCCCCCCTCCCCACCCCCAATTTTGTATTTATTTATTTTTTAATTATTTTGTGCAGCGATGGGGGCGGGGGGGGGGGGGGGGCGCGCGCCAGGCGGGGCGGGGCGGGGCGAGGGGCGGGGCGGGGCGAGGCGGAGAGGTGCGGCGGCAGCCAATCAGAGCGGCGCGCTCCGAAAGTTTCCTTTTATGGCGAGGCGGCGGCGGCGGCGGCCCTATAAAAAGCGAAGCGCGCGGCGGGCGGGAGTCGCTGCGCGCTGCCTTCGCCCCGTGCCCCGCTCCGCCGCCGCCTCGCGCCGCCCGCCCCGGCTCTGACTGACCGCGTTACTCCCACAGGTGAGCGGGCGGGACGGCCCTTCTCCTCCGGGCTGTAATTAGCTGAGCAAGAGGTAAGGGTTTAAGGGATGGTTGGTTGGTGGGGTATTAATGTTTAATTACCTGGAGCACCTGCCTGAAATCACTTTTTTTCAGGTTGGACCGGTGCCACCATGGACTATAAGGACCACGACGGAGACTACAAGGATCATGATATTGATTACAAAGACGATGACGATAAGATGGCCCCAAAGAAGAAGCGGAAGGTCGGTATCCACGGAGTCCCAGCAGCCGACAAGAAGTACAGCATCGGCCTGGACATCGGCACCAACTCTGTGGGCTGGGCCGTGATCACCGACGAGTACAAGGTGCCCAGCAAGAAATTCAAGGTGCTGGGCAACACCGACCGGCACAGCATCAAGAAGAACCTGATCGGAGCCCTGCTGTTCGACAGCGGCGAAACAGCCGAGGCCACCCGGCTGAAGAGAACCGCCAGAAGAAGATACACCAGACGGAAGAACCGGATCTGCTATCTGCAAGAGATCTTCAGCAACGAGATGGCCAAGGTGGACGACAGCTTCTTCCACAGACTGGAAGAGTCCTTCCTGGTGGAAGAGGATAAGAAGCACGAGCGGCACCCCATCTTCGGCAACATCGTGGACGAGGTGGCCTACCACGAGAAGTACCCCACCATCTACCACCTGAGAAAGAAACTGGTGGACAGCACCGACAAGGCCGACCTGCGGCTGATCTATCTGGCCCTGGCCCACATGATCAAGTTCCGGGGCCACTTCCTGATCGAGGGCGACCTGAACCCCGACAACAGCGACGTGGACAAGCTGTTCATCCAGCTGGTGCAGACCTACAACCAGCTGTTCGAGGAAAACCCCATCAACGCCAGCGGCGTGGACGCCAAGGCCATCCTGTCTGCCAGACTGAGCAAGAGCAGACGGCTGGAAAATCTGATCGCCCAGCTGCCCGGCGAGAAGAAGAATGGCCTGTTCGGAAACCTGATTGCCCTGAGCCTGGGCCTGACCCCCAACTTCAAGAGCAACTTCGACCTGGCCGAGGATGCCAAACTGCAGCTGAGCAAGGACACCTACGACGACGACCTGGACAACCTGCTGGCCCAGATCGGCGACCAGTACGCCGACCTGTTTCTGGCCGCCAAGAACCTGTCCGACGCCATCCTGCTGAGCGACATCCTGAGAGTGAACACCGAGATCACCAAGGCCCCCCTGAGCGCCTCTATGATCAAGAGATACGACGAGCACCACCAGGACCTGACCCTGCTGAAAGCTCTCGTGCGGCAGCAGCTGCCTGAGAAGTACAAAGAGATTTTCTTCGACCAGAGCAAGAACGGCTACGCCGGCTACATTGACGGCGGAGCCAGCCAGGAAGAGTTCTACAAGTTCATCAAGCCCATCCTGGAAAAGATGGACGGCACCGAGGAACTGCTCGTGAAGCTGAACAGAGAGGACCTGCTGCGGAAGCAGCGGACCTTCGACAACGGCAGCATCCCCCACCAGATCCACCTGGGAGAGCTGCACGCCATTCTGCGGCGGCAGGAAGATTTTTACCCATTCCTGAAGGACAACCGGGAAAAGATCGAGAAGATCCTGACCTTCCGCATCCCCTACTACGTGGGCCCTCTGGCCAGGGGAAACAGCAGATTCGCCTGGATGACCAGAAAGAGCGAGGAAACCATCACCCCCTGGAACTTCGAGGAAGTGGTGGACAAGGGCGCTTCCGCCCAGAGCTTCATCGAGCGGATGACCAACTTCGATAAGAACCTGCCCAACGAGAAGGTGCTGCCCAAGCACAGCCTGCTGTACGAGTACTTCACCGTGTATAACGAGCTGACCAAAGTGAAATACGTGACCGAGGGAATGAGAAAGCCCGCCTTCCTGAGCGGCGAGCAGAAAAAGGCCATCGTGGACCTGCTGTTCAAGACCAACCGGAAAGTGACCGTGAAGCAGCTGAAAGAGGACTACTTCAAGAAAATCGAGTGCTTCGACTCCGTGGAAATCTCCGGCGTGGAAGATCGGTTCAACGCCTCCCTGGGCACATACCACGATCTGCTGAAAATTATCAAGGACAAGGACTTCCTGGACAATGAGGAAAACGAGGACATTCTGGAAGATATCGTGCTGACCCTGACACTGTTTGAGGACAGAGAGATGATCGAGGAACGGCTGAAAACCTATGCCCACCTGTTCGACGACAAAGTGATGAAGCAGCTGAAGCGGCGGAGATACACCGGCTGGGGCAGGCTGAGCCGGAAGCTGATCAACGGCATCCGGGACAAGCAGTCCGGCAAGACAATCCTGGATTTCCTGAAGTCCGACGGCTTCGCCAACAGAAACTTCATGCAGCTGATCCACGACGACAGCCTGACCTTTAAAGAGGACATCCAGAAAGCCCAGGTGTCCGGCCAGGGCGATAGCCTGCACGAGCACATTGCCAATCTGGCCGGCAGCCCCGCCATTAAGAAGGGCATCCTGCAGACAGTGAAGGTGGTGGACGAGCTCGTGAAAGTGATGGGCCGGCACAAGCCCGAGAACATCGTGATCGAAATGGCCAGAGAGAACCAGACCACCCAGAAGGGACAGAAGAACAGCCGCGAGAGAATGAAGCGGATCGAAGAGGGCATCAAAGAGCTGGGCAGCCAGATCCTGAAAGAACACCCCGTGGAAAACACCCAGCTGCAGAACGAGAAGCTGTACCTGTACTACCTGCAGAATGGGCGGGATATGTACGTGGACCAGGAACTGGACATCAACCGGCTGTCCGACTACGATGTGGACCATATCGTGCCTCAGAGCTTTCTGAAGGACGACTCCATCGACAACAAGGTGCTGACCAGAAGCGACAAGAACCGGGGCAAGAGCGACAACGTGCCCTCCGAAGAGGTCGTGAAGAAGATGAAGAACTACTGGCGGCAGCTGCTGAACGCCAAGCTGATTACCCAGAGAAAGTTCGACAATCTGACCAAGGCCGAGAGAGGCGGCCTGAGCGAACTGGATAAGGCCGGCTTCATCAAGAGACAGCTGGTGGAAACCCGGCAGATCACAAAGCACGTGGCACAGATCCTGGACTCCCGGATGAACACTAAGTACGACGAGAATGACAAGCTGATCCGGGAAGTGAAAGTGATCACCCTGAAGTCCAAGCTGGTGTCCGATTTCCGGAAGGATTTCCAGTTTTACAAAGTGCGCGAGATCAACAACTACCACCACGCCCACGACGCCTACCTGAACGCCGTCGTGGGAACCGCCCTGATCAAAAAGTACCCTAAGCTGGAAAGCGAGTTCGTGTACGGCGACTACAAGGTGTACGACGTGCGGAAGATGATCGCCAAGAGCGAGCAGGAAATCGGCAAGGCTACCGCCAAGTACTTCTTCTACAGCAACATCATGAACTTTTTCAAGACCGAGATTACCCTGGCCAACGGCGAGATCCGGAAGCGGCCTCTGATCGAGACAAACGGCGAAACCGGGGAGATCGTGTGGGATAAGGGCCGGGATTTTGCCACCGTGCGGAAAGTGCTGAGCATGCCCCAAGTGAATATCGTGAAAAAGACCGAGGTGCAGACAGGCGGCTTCAGCAAAGAGTCTATCCTGCCCAAGAGGAACAGCGATAAGCTGATCGCCAGAAAGAAGGACTGGGACCCTAAGAAGTACGGCGGCTTCGACAGCCCCACCGTGGCCTATTCTGTGCTGGTGGTGGCCAAAGTGGAAAAGGGCAAGTCCAAGAAACTGAAGAGTGTGAAAGAGCTGCTGGGGATCACCATCATGGAAAGAAGCAGCTTCGAGAAGAATCCCATCGACTTTCTGGAAGCCAAGGGCTACAAAGAAGTGAAAAAGGACCTGATCATCAAGCTGCCTAAGTACTCCCTGTTCGAGCTGGAAAACGGCCGGAAGAGAATGCTGGCCTCTGCCGGCGAACTGCAGAAGGGAAACGAACTGGCCCTGCCCTCCAAATATGTGAACTTCCTGTACCTGGCCAGCCACTATGAGAAGCTGAAGGGCTCCCCCGAGGATAATGAGCAGAAACAGCTGTTTGTGGAACAGCACAAGCACTACCTGGACGAGATCATCGAGCAGATCAGCGAGTTCTCCAAGAGAGTGATCCTGGCCGACGCTAATCTGGACAAAGTGCTGTCCGCCTACAACAAGCACCGGGATAAGCCCATCAGAGAGCAGGCCGAGAATATCATCCACCTGTTTACCCTGACCAATCTGGGAGCCCCTGCCGCCTTCAAGTACTTTGACACCACCATCGACCGGAAGAGGTACACCAGCACCAAAGAGGTGCTGGACGCCACCCTGATCCACCAGAGCATCACCGGCCTGTACGAGACACGGATCGACCTGTCTCAGCTGGGAGGCGACAAAAGGCCGGCGGCCACGAAAAAGGCCGGCCAGGCAAAAAAGAAAAAGGAATTCGGCAGTGGAGGATCCGGCGCAACAAACTTCTCTCTGCTGAAACAAGCCGGAGATGTCGAAGAGAATCCTGGACCGTCCGAGGCACCCCGGGCCGAGACCTTTGTGTTCCTGGACCTGGAAGCCACTGGGCTCCCCAGTGTGGAGCCCGAGATTGCCGAGCTGTCCCTCTTTGCTGTCCACCGCTCCTCCCTGGAGAACCCGGAGCACGACGAGTCTGGTGCCCTAGTATTGCCCCGGGTCCTGGACAAGCTCACGCTGTGCATGTGCCCGGAGCGCCCCTTCACTGCCAAGGCCAGCGAGATCACCGGCCTGAGCAGTGAGGGCCTGGCGCGATGCCGGAAGGCTGGCTTTGATGGCGCCGTGGTGCGGACGCTGCAGGCCTTCCTGAGCCGCCAGGCAGGGCCCATCTGCCTTGTGGCCCACAATGGCTTTGATTATGATTTCCCCCTGCTGTGTGCCGAGCTGCGGCGCCTGGGTGCTCGCCTGCCTCGGGACACTGTCTGCCTGGACACGCTGCCAGCCCTGCGGGGCCTGGACCGCGCCCACAGCCACGGCACACGAGCTAGAGGCAGACAGGGTTACAGCCTCGGCAGCCTCTTCCACCGCTACTTCCGGGCAGAGCCAAGCGCAGCCCACTCAGCCGAGGGCGACGTGCACACCCTGCTCCTGATCTTCCTGCACCGCGCCGCAGAGCTGCTCGCCTGGGCCGATGAGCAGGCCCGTGGGTGGGCCCACATCGAGCCCATGTACTTGCCGCCTGATGACCCCAGCCTGGAGGCCGAGGGCAGAGGAAGTCTGCTAACATGCGGTGACGTCGAGGAGAATCCTGGCCCAGTGAGCAAGGGCGAGGAGCTGTTCACCGGGGTGGTGCCCATCCTGGTCGAGCTGGACGGCGACGTAAACGGCCACAAGTTCAGCGTGTCCGGCGAGGGCGAGGGCGATGCCACCTACGGCAAGCTGACCCTGAAGTTCATCTGCACCACCGGCAAGCTGCCCGTGCCCTGGCCCACCCTCGTGACCACCCTGACCTACGGCGTGCAGTGCTTCAGCCGCTACCCCGACCACATGAAGCAGCACGACTTCTTCAAGTCCGCCATGCCCGAAGGCTACGTCCAGGAGCGCACCATCTTCTTCAAGGACGACGGCAACTACAAGACCCGCGCCGAGGTGAAGTTCGAGGGCGACACCCTGGTGAACCGCATCGAGCTGAAGGGCATCGACTTCAAGGAGGACGGCAACATCCTGGGGCACAAGCTGGAGTACAACTACAACAGCCACAACGTCTATATCATGGCCGACAAGCAGAAGAACGGCATCAAGGTGAACTTCAAGATCCGCCACAACATCGAGGACGGCAGCGTGCAGCTCGCCGACCACTACCAGCAGAACACCCCCATCGGCGACGGCCCCGTGCTGCTGCCCGACAACCACTACCTGAGCACCCAGTCCGCCCTGAGCAAAGACCCCAACGAGAAGCGCGATCACATGGTCCTGCTGGAGTTCGTGACCGCCGCCGGGATCACTCTCGGCATGGACGAGCTGTACAAGGAATTCTAACTAGAGCTCGCTGATCAGCCTCGACTGTGCCTTCTAGTTGCCAGCCATCTGTTGTTTGCCCCTCCCCCGTGCCTTCCTTGACCCTGGAAGGTGCCACTCCCACTGTCCTTTCCTAATAAAATGAGGAAATTGCATCGCATTGTCTGAGTAGGTGTCATTCTATTCTGGGGGGTGGGGTGGGGCAGGACAGCAAGGGGGAGGATTGGGAAGAGAATAGCAGGCATGCTGGGGAGCGGCCGCAGGAACCCCTAGTGATGGAGTTGGCCACTCCCTCTCTGCGCGCTCGCTCGCTCACTGAGGCCGGGCGACCAAAGGTCGCCCGACGCCCGGGCTTTGCCCGGGCGGCCTCAGTGAGCGAGCGAGCGCGCAGCTGCCTGCAGGGGCGCCTGATGCGGTATTTTCTCCTTACGCATCTGTGCGGTATTTCACACCGCATACGTCAAAGCAACCATAGTACGCGCCCTGTAGCGGCGCATTAAGCGCGGCGGGTGTGGTGGTTACGCGCAGCGTGACCGCTACACTTGCCAGCGCCTTAGCGCCCGCTCCTTTCGCTTTCTTCCCTTCCTTTCTCGCCACGTTCGCCGGCTTTCCCCGTCAAGCTCTAAATCGGGGGCTCCCTTTAGGGTTCCGATTTAGTGCTTTACGGCACCTCGACCCCAAAAAACTTGATTTGGGTGATGGTTCACGTAGTGGGCCATCGCCCTGATAGACGGTTTTTCGCCCTTTGACGTTGGAGTCCACGTTCTTTAATAGTGGACTCTTGTTCCAAACTGGAACAACACTCAACTCTATCTCGGGCTATTCTTTTGATTTATAAGGGATTTTGCCGATTTCGGTCTATTGGTTAAAAAATGAGCTGATTTAACAAAAATTTAACGCGAATTTTAACAAAATATTAACGTTTACAATTTTATGGTGCACTCTCAGTACAATCTGCTCTGATGCCGCATAGTTAAGCCAGCCCCGACACCCGCCAACACCCGCTGACGCGCCCTGACGGGCTTGTCTGCTCCCGGCATCCGCTTACAGACAAGCTGTGACCGTCTCCGGGAGCTGCATGTGTCAGAGGTTTTCACCGTCATCACCGAAACGCGCGAGACGAAAGGGCCTCGTGATACGCCTATTTTTATAGGTTAATGTCATGATAATAATGGTTTCTTAGACGTCAGGTGGCACTTTTCGGGGAAATGTGCGCGGAACCCCTATTTGTTTATTTTTCTAAATACATTCAAATATGTATCCGCTCATGAGACAATAACCCTGATAAATGCTTCAATAATATTGAAAAAGGAAGAGTATGAGTATTCAACATTTCCGTGTCGCCCTTATTCCCTTTTTTGCGGCATTTTGCCTTCCTGTTTTTGCTCACCCAGAAACGCTGGTGAAAGTAAAAGATGCTGAAGATCAGTTGGGTGCACGAGTGGGTTACATCGAACTGGATCTCAACAGCGGTAAGATCCTTGAGAGTTTTCGCCCCGAAGAACGTTTTCCAATGATGAGCACTTTTAAAGTTCTGCTATGTGGCGCGGTATTATCCCGTATTGACGCCGGGCAAGAGCAACTCGGTCGCCGCATACACTATTCTCAGAATGACTTGGTTGAGTACTCACCAGTCACAGAAAAGCATCTTACGGATGGCATGACAGTAAGAGAATTATGCAGTGCTGCCATAACCATGAGTGATAACACTGCGGCCAACTTACTTCTGACAACGATCGGAGGACCGAAGGAGCTAACCGCTTTTTTGCACAACATGGGGGATCATGTAACTCGCCTTGATCGTTGGGAACCGGAGCTGAATGAAGCCATACCAAACGACGAGCGTGACACCACGATGCCTGTAGCAATGGCAACAACGTTGCGCAAACTATTAACTGGCGAACTACTTACTCTAGCTTCCCGGCAACAATTAATAGACTGGATGGAGGCGGATAAAGTTGCAGGACCACTTCTGCGCTCGGCCCTTCCGGCTGGCTGGTTTATTGCTGATAAATCTGGAGCCGGTGAGCGTGGAAGCCGCGGTATCATTGCAGCACTGGGGCCAGATGGTAAGCCCTCCCGTATCGTAGTTATCTACACGACGGGGAGTCAGGCAACTATGGATGAACGAAATAGACAGATCGCTGAGATAGGTGCCTCACTGATTAAGCATTGGTAACTGTCAGACCAAGTTTACTCATATATACTTTAGATTGATTTAAAACTTCATTTTTAATTTAAAAGGATCTAGGTGAAGATCCTTTTTGATAATCTCATGACCAAAATCCCTTAACGTGAGTTTTCGTTCCACTGAGCGTCAGACCCCGTAGAAAAGATCAAAGGATCTTCTTGAGATCCTTTTTTTCTGCGCGTAATCTGCTGCTTGCAAACAAAAAAACCACCGCTACCAGCGGTGGTTTGTTTGCCGGATCAAGAGCTACCAACTCTTTTTCCGAAGGTAACTGGCTTCAGCAGAGCGCAGATACCAAATACTGTTCTTCTAGTGTAGCCGTAGTTAGGCCACCACTTCAAGAACTCTGTAGCACCGCCTACATACCTCGCTCTGCTAATCCTGTTACCAGTGGCTGCTGCCAGTGGCGATAAGTCGTGTCTTACCGGGTTGGACTCAAGACGATAGTTACCGGATAAGGCGCAGCGGTCGGGCTGAACGGGGGGTTCGTGCACACAGCCCAGCTTGGAGCGAACGACCTACACCGAACTGAGATACCTACAGCGTGAGCTATGAGAAAGCGCCACGCTTCCCGAAGGGAGAAAGGCGGACAGGTATCCGGTAAGCGGCAGGGTCGGAACAGGAGAGCGCACGAGGGAGCTTCCAGGGGGAAACGCCTGGTATCTTTATAGTCCTGTCGGGTTTCGCCACCTCTGACTTGAGCGTCGATTTTTGTGATGCTCGTCAGGGGGGCGGAGCCTATGGAAAAACGCCAGCAACGCGGCCTTTTTACGGTTCCTGGCCTTTTGCTGGCCTTTTGCTCACATGT

Legend: U6 Promoter, gRNA scaffold, CAG promoter, TREX2, SpCas9, NLS signals, P2A signal, T2A signal, EGFP

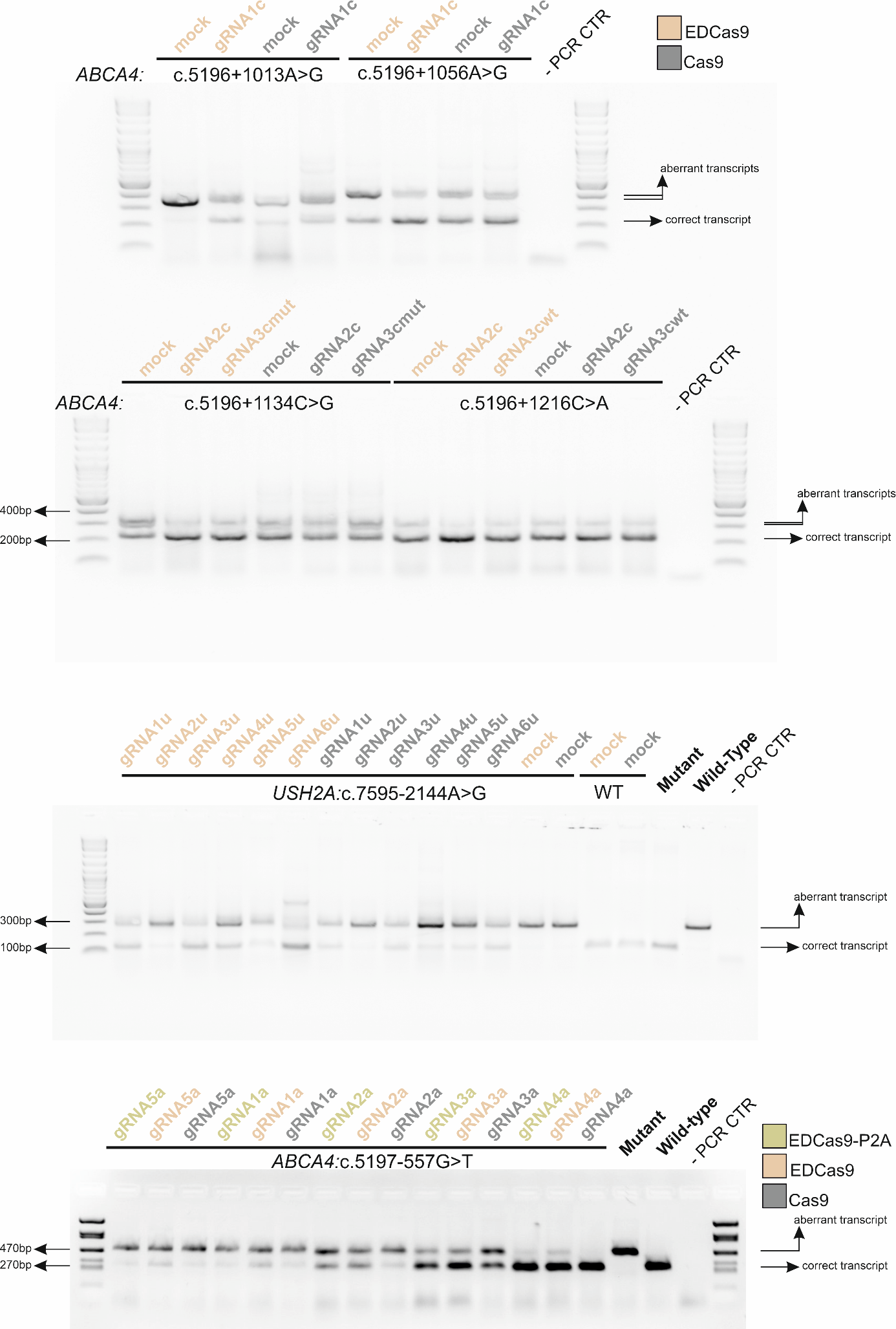

**Supplementary Figure 1: Representative agarose gel picture showing splicing rescue in minigene assay for the isolated ABCA4:c.5197-557G>T deep-intronic variant.** Co-transfection of the mutant minigene with the selected five single gRNAs (gRNA1a, gRNA2a, gRNA3a, gRNA4a, and gRNA5a) paired to EDCas9-T2A (NLS-TREX2-T2A-SpCas9-NLS), resulting in the independent translation of TREX2 and SpCas9, EDCas9 (NLS-TREX2-linker-SpCas9-NLS), the chimeric fusion protein of TREX2 to SpCas9, or the wild-type SpCas9 (Cas9) orthologue. PCR control without DNA template (-PCR CTR).

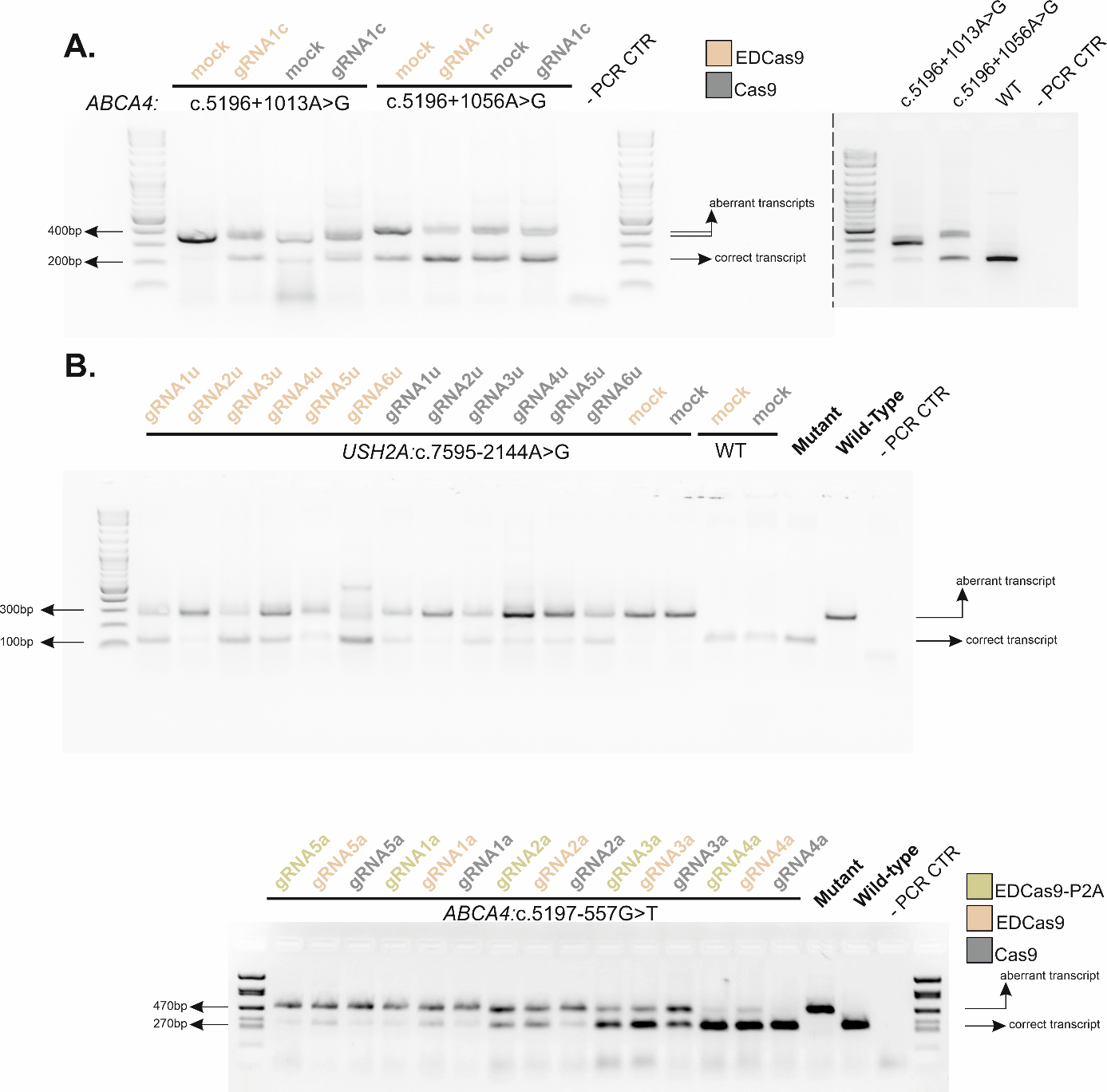

**Supplementary Figure 2: Representative agarose gel picture showing splicing rescue in minigene assay for the clustered deep-intronic variants ABCA4:c.5196+1013A>G and ABCA4:c.5196+1056A>G and the common isolated deep-intronic variant USH2A:c.7595-2144A>G.** Co-transfection of mutant minigenes with the different single gRNAs paired to EDCas9 and SpCas9 (Cas9). **A)** The selected clustered ABCA4:c.5196+1013A>G and ABCA4:c.5196+1056A>G were addressed by single gRNA1c. **B**) the common USH2A:c.7595-2144A>G deep-intronic variant was addressed by six gRNAs (gRNA1u, gRNA2u, gRNA3u, gRNA4u, gRNA5u, and gRNA6u). A mock gRNA (mock) coupled to EDCas9 and Cas9 was used as a control. Only mutant and WT minigene plasmids and PCR control without DNA template (-PCR CTR) were used as controls The dashed vertical lines separate two agarose gel pictures.

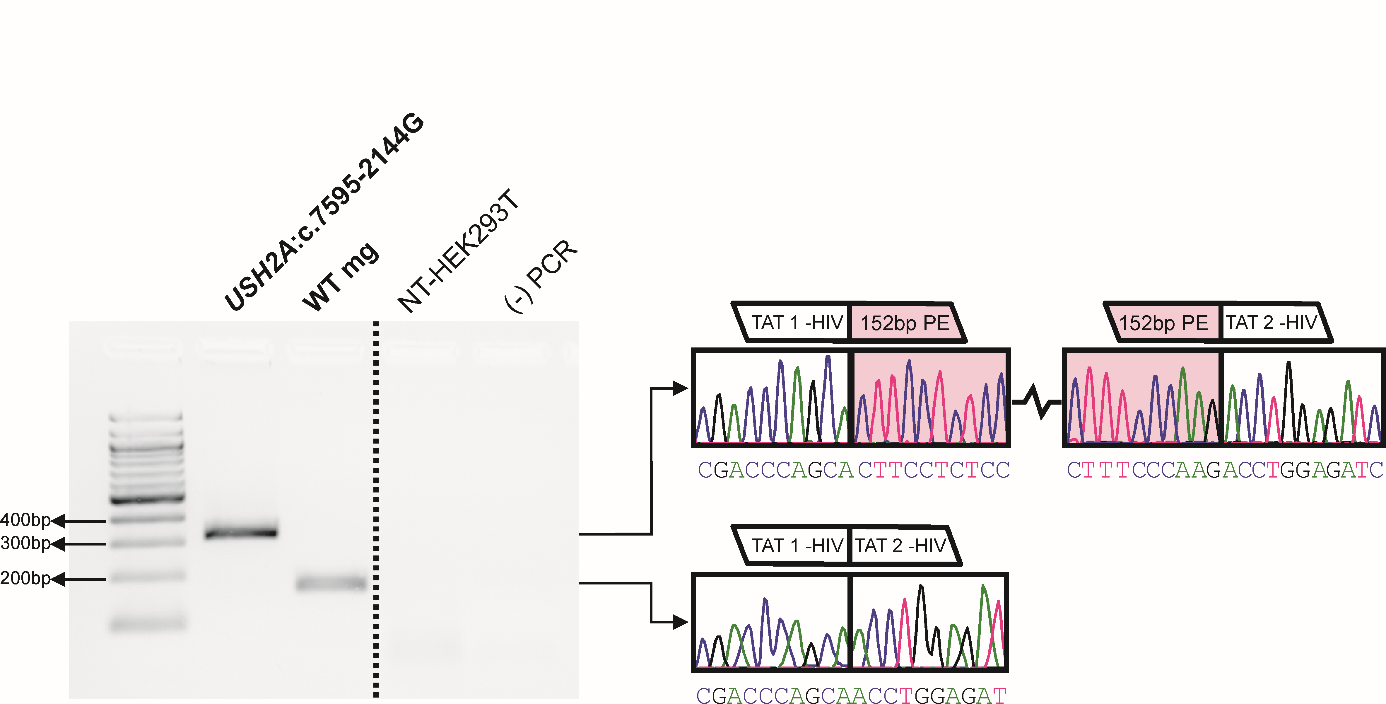

**Supplementary Figure 3: Minigene splicing pattern for the USH2A:c.7595-2144A>G DIV in HEK293T.** The agarose gel displays amplicon bands from reverse transcription PCR analysis of mutant USH2A (USH2A:c.7595-2144G) and wild-type minigene (mg) plasmids. Non-transfected HEK293T (NT-HEK293T) and the no DNA (-) PCR are shown as controls. PCR amplicon sequencing confirms the presence of the 152 bp pseudoexon (PE) sequence in the mutant USH2A and the correctly spliced cDNA sequence in the wild-type minigene.

.
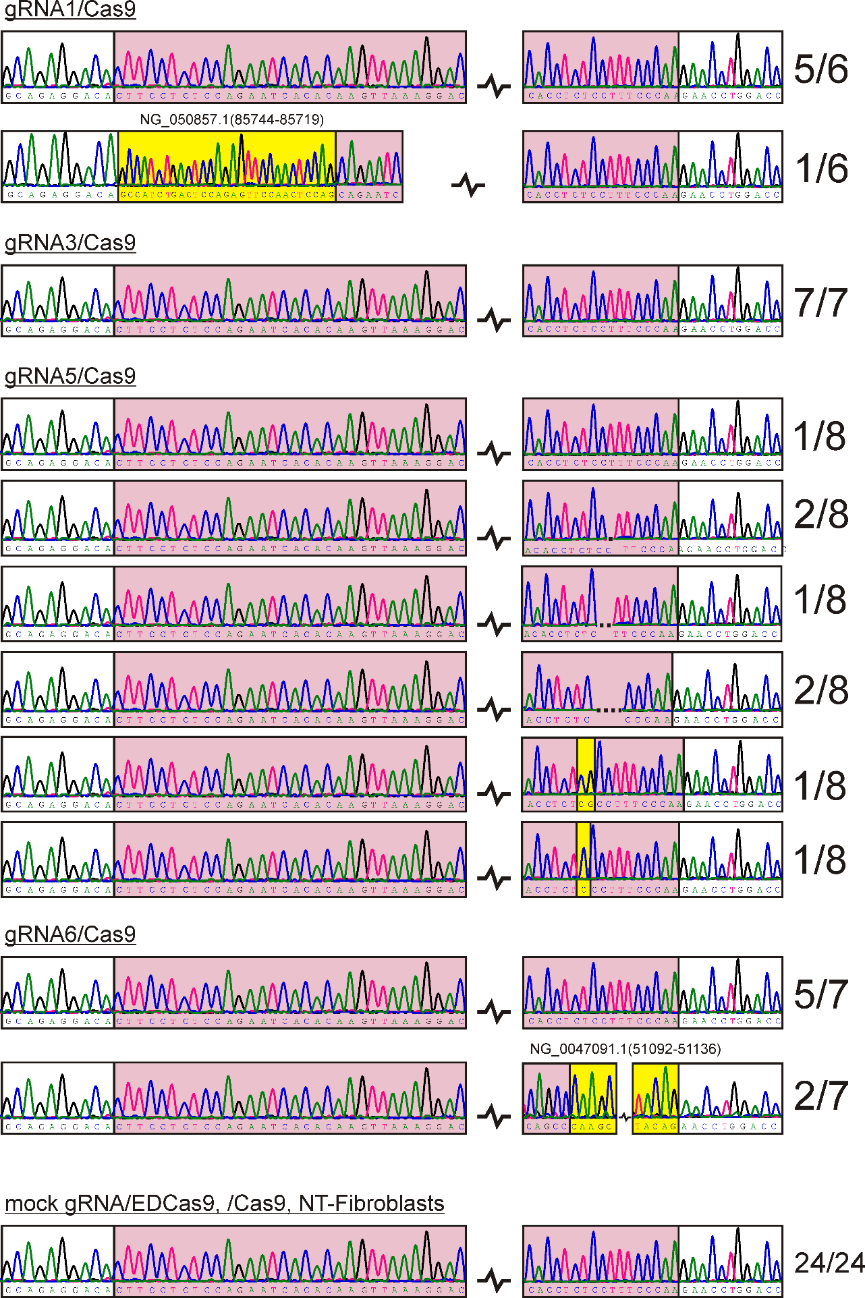

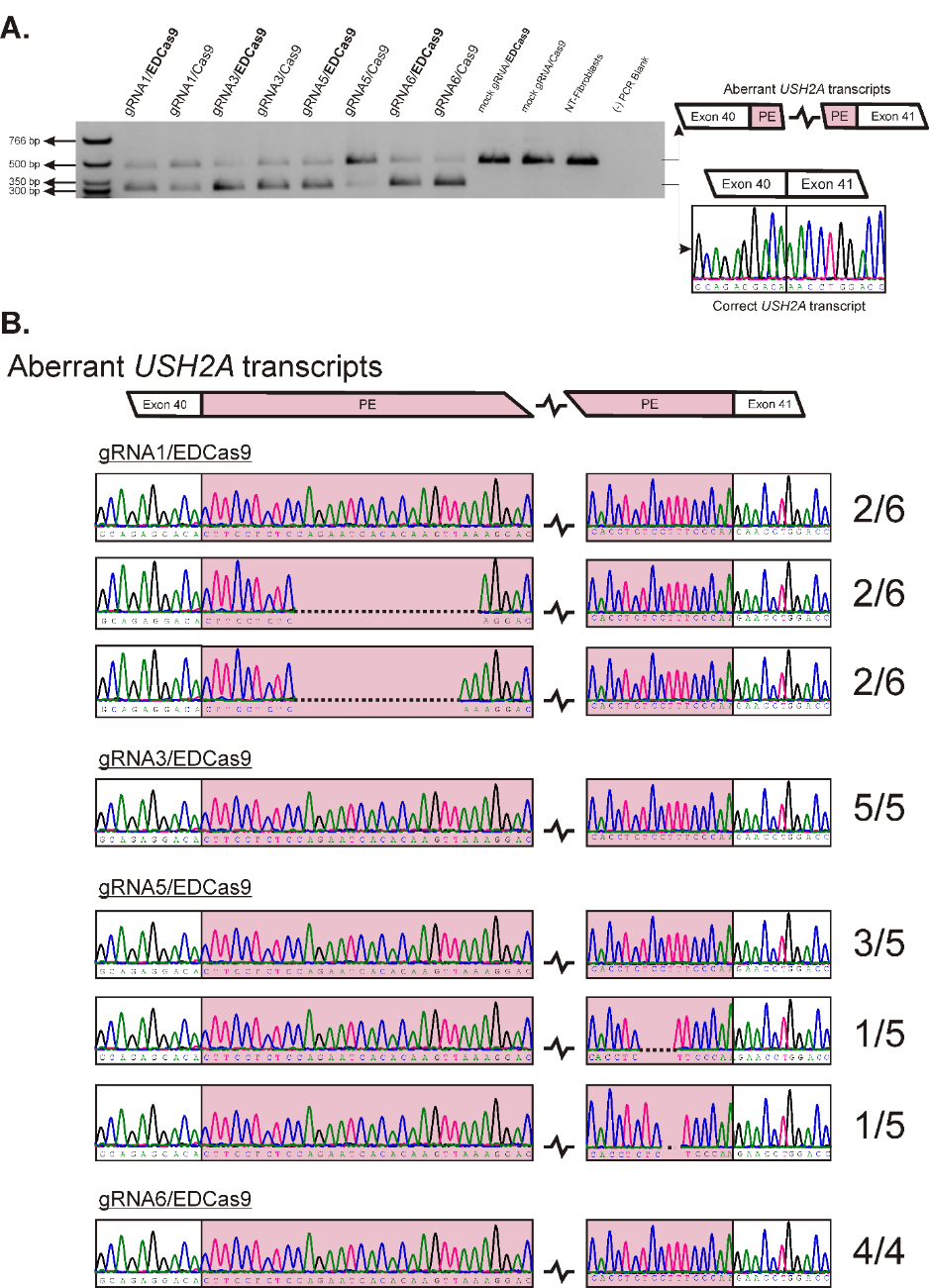

**Supplementary Figure 4: Characterization of the splicing pattern in homozygous USH2A:c.7595-2144G patient-derived fibroblasts upon EDCas9 and Cas9-mediated splicing correction.**
(**A**) Representative agarose gel showing the splicing pattern upon EDCas9 and Cas9 editing. The aberrant transcripts are represented by the top band, while the rescued USH2A transcript is represented by the bottom band. (**B**) Sequencing characterization of aberrant transcripts upon EDCas9 and Cas9 editing. Individual sub-cloned transcripts were sequenced and aligned to the reference aberrant USH2A transcript sequence containing the 152 bp pseudoexon (PE). Some USH2A aberrant transcripts show shorter pseudoexon inclusion (dashed lines) upon EDCas9 and Cas9 editing. Single or double insertion of nucleotides (in yellow) retained in the aberrant transcripts are evident for gRNA5/Cas9. Insertion of larger sequence stretches (in yellow) mapped on different genes were detected for gRNA1/Cas9 and gRNA6/Cas9. The ratio of characterized sequence is reported on the right end side of each electropherogram.

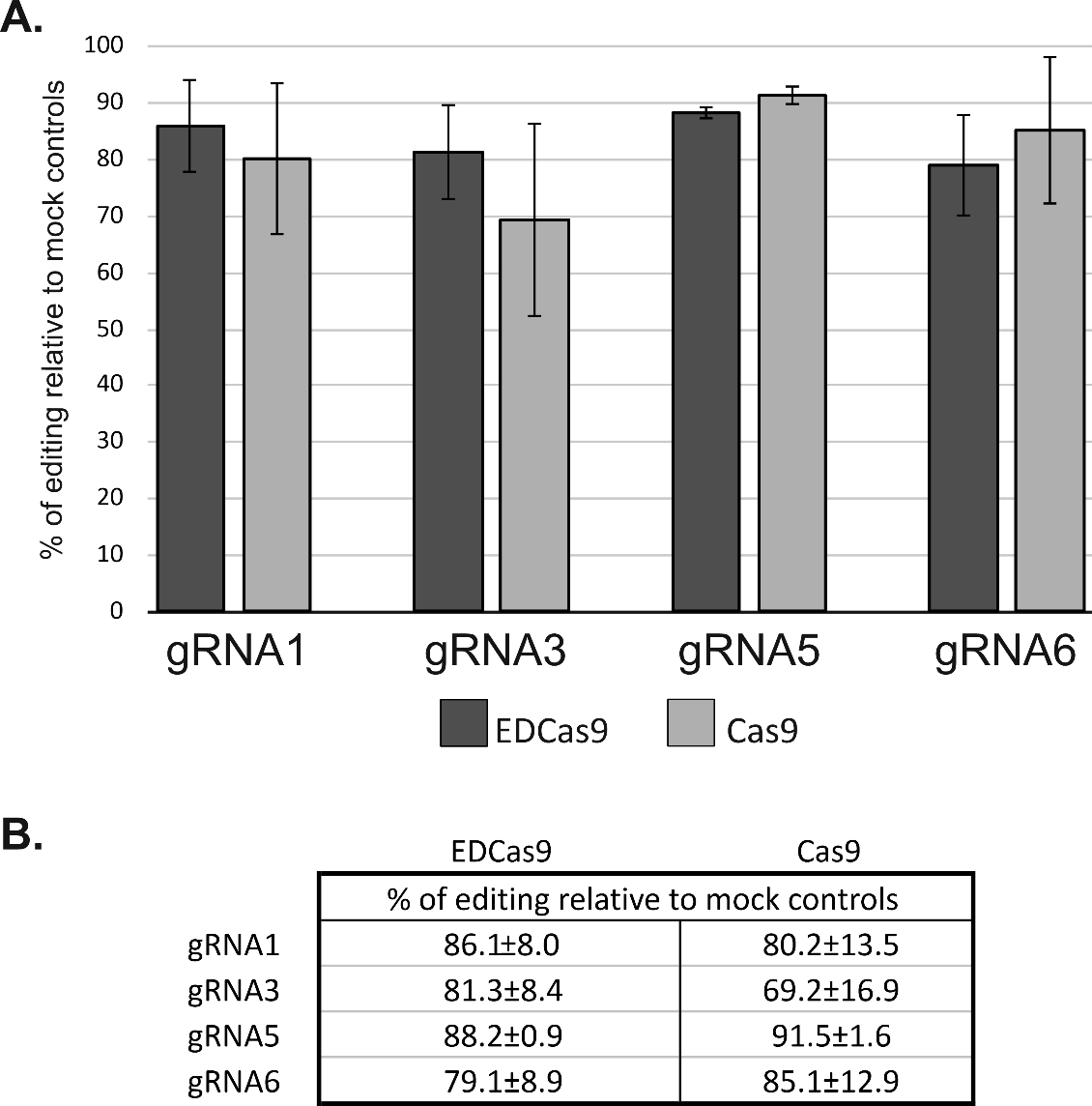

**Supplementary Figure 5**: **Gene editing efficiency in homozygous USH2A:c.7595-2144G patient-derived fibroblasts. (A)** Bar graph showing the percentage of gene editing normalized to mock controls. (B) Table reporting the relative values for the % of editing relative to mock controls. Results are presented as mean (%) ± SD of n=3 biological replicates. Difference between EDCas9 and Cas9 is **NOT** significant for any gRNA.

**gRNA1/EDCas9**

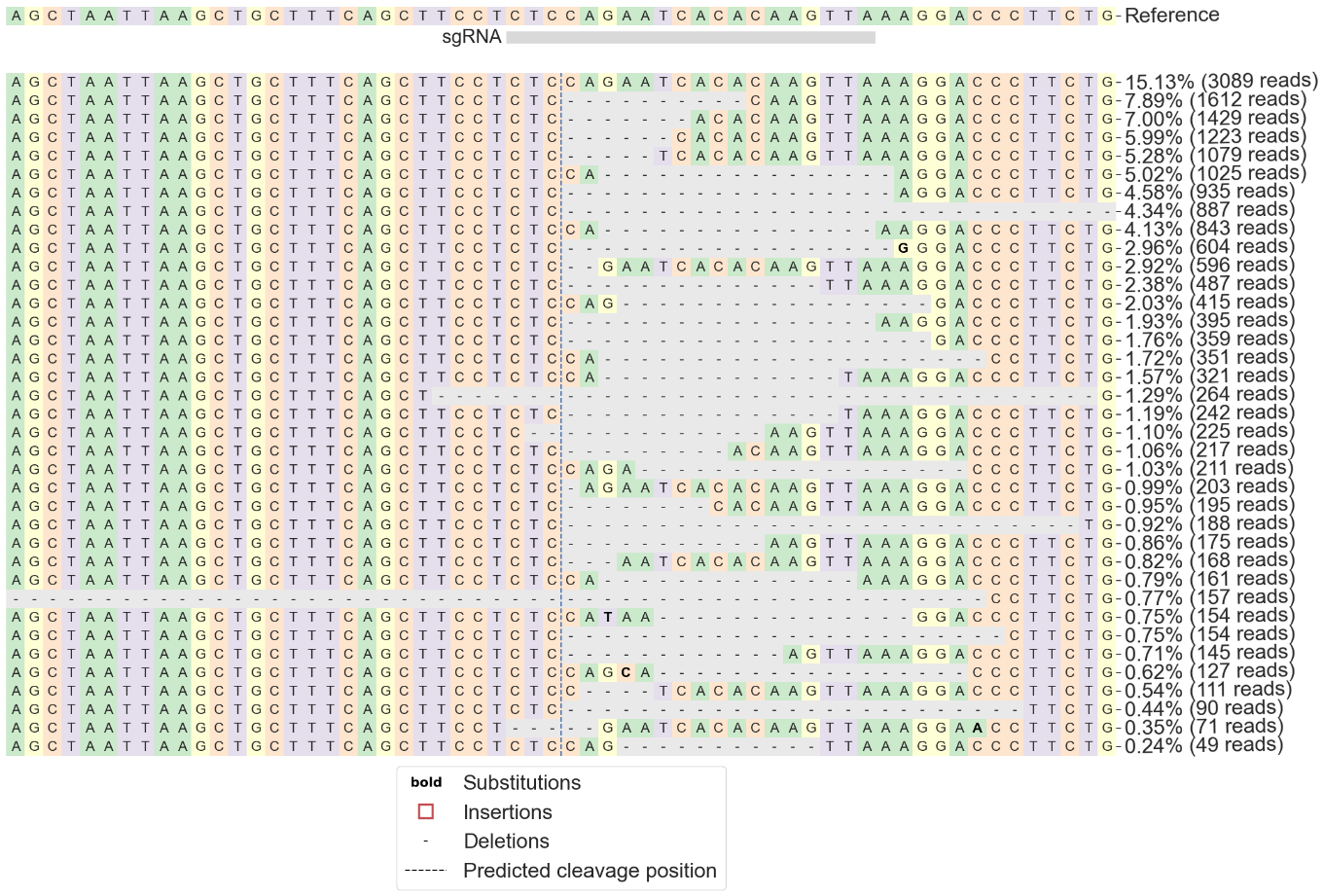

**gRNA3/EDCas9**

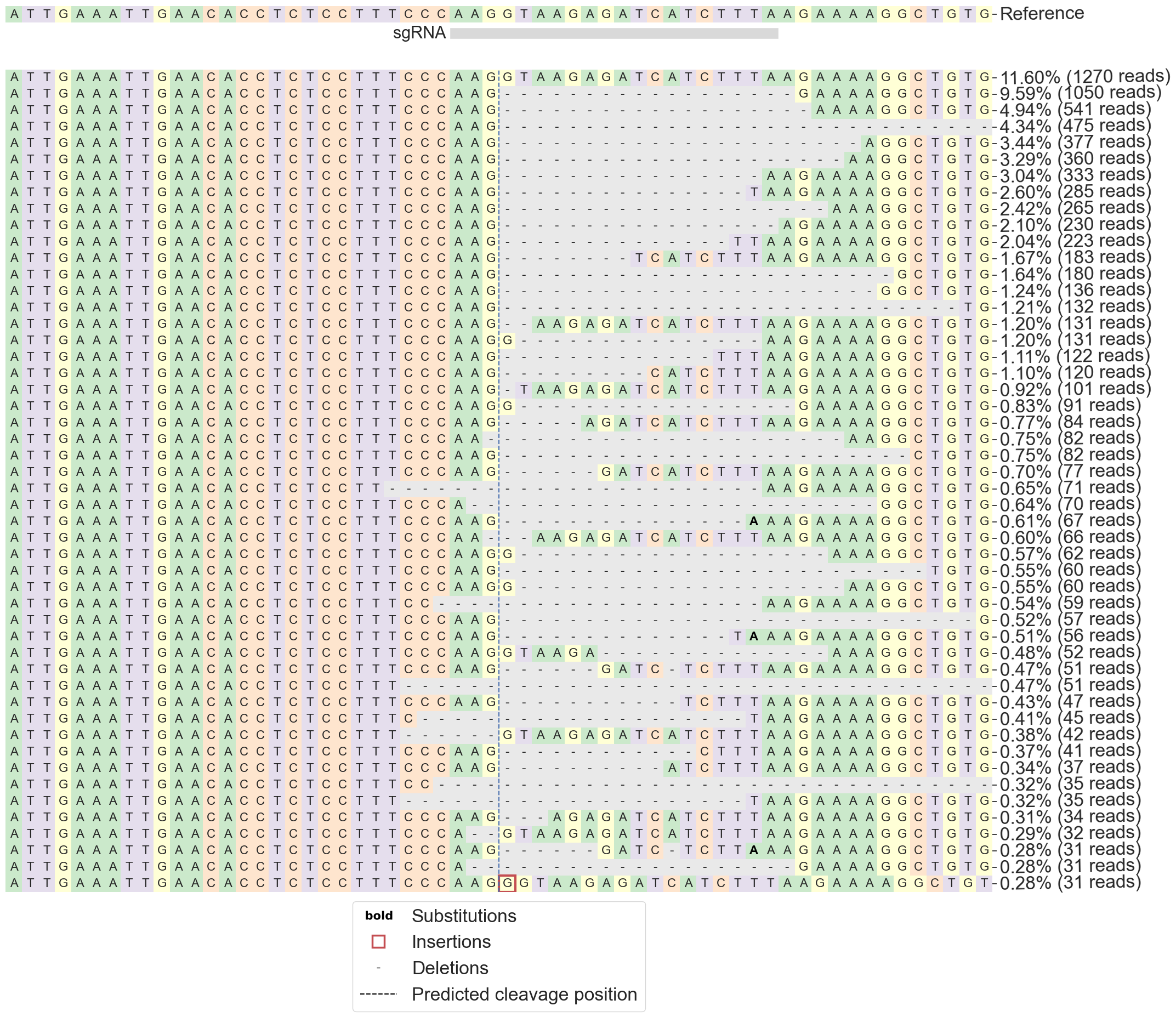

**gRNA5/EDCas9**

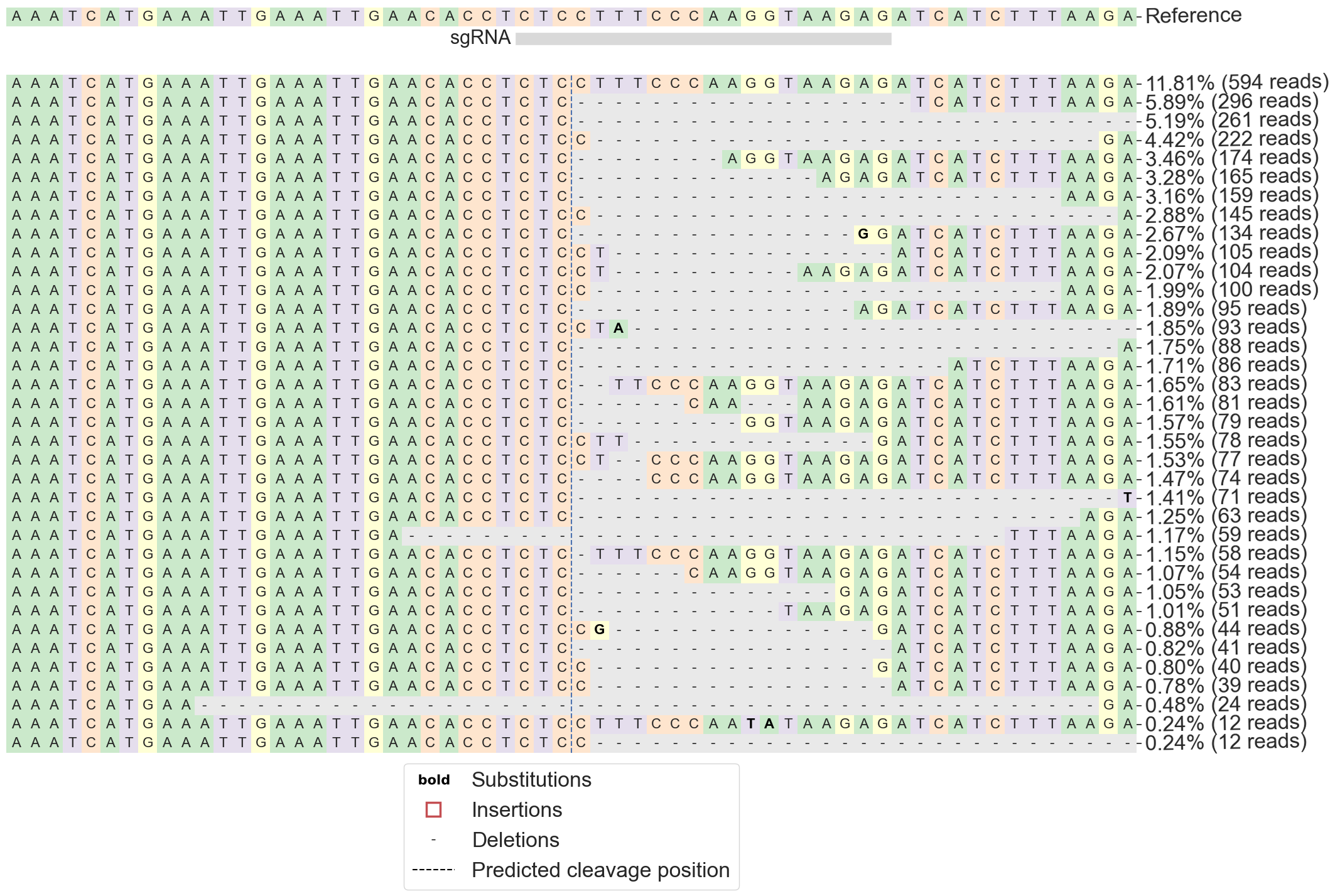

**gRNA6/EDCas9**

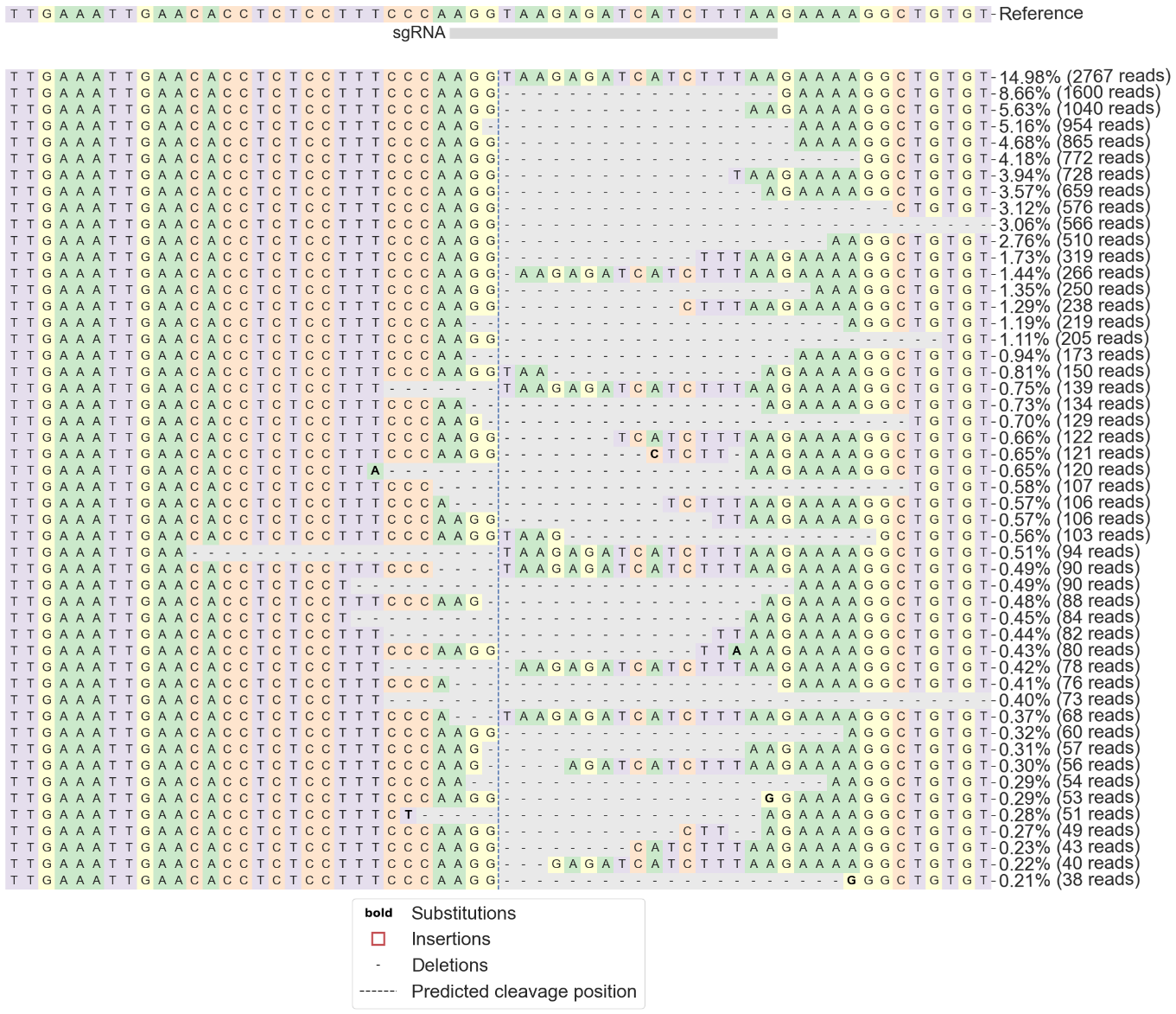

**gRNA6/Cas9**

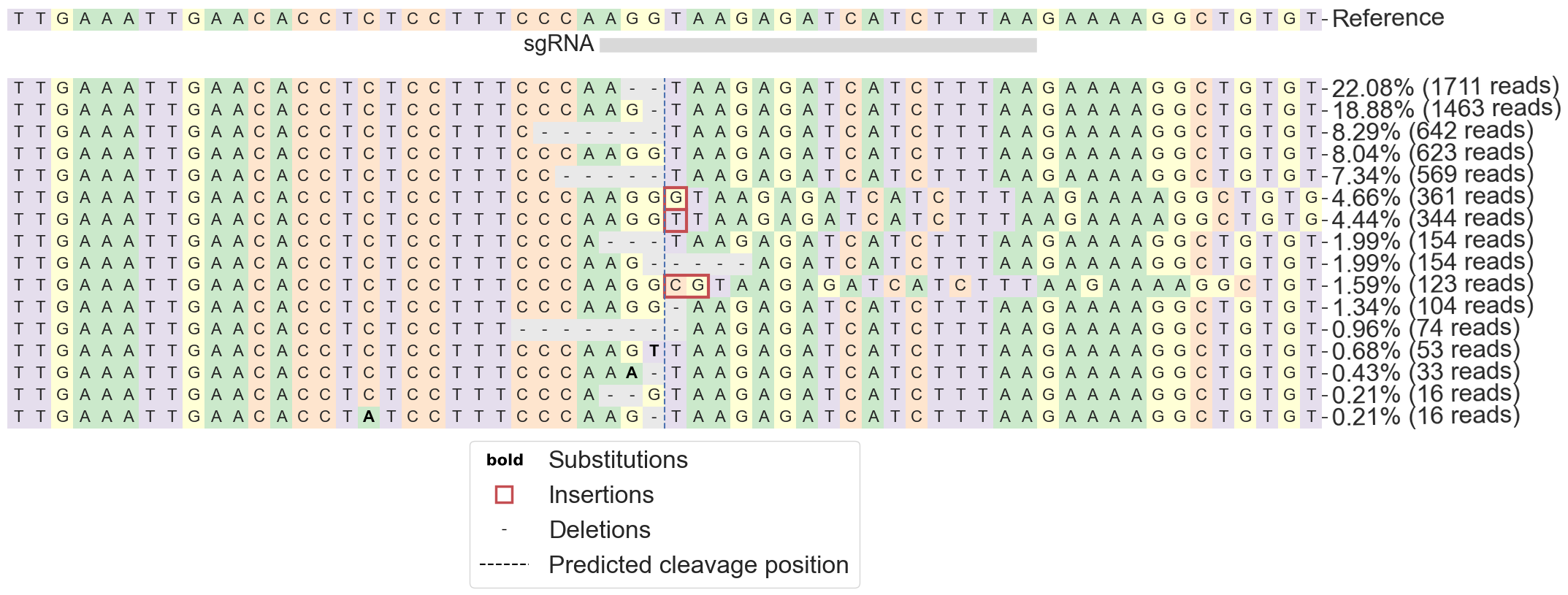

**Supplementary Figure 6: Representative examples of the directional deletion profiles induced by EDCas9 for gRNA1, 3, 5 and 6 in homozygous USH2A:c.7595-2144GG patient-derived fibroblasts and gRNA6/Cas9.** The CRISPResso results correspond to one biological replicate selected as representative for each gRNA/EDCas9 or gRNA6/Cas9 combination, with the cut site marked by a dashed line. The figures display the 10 most common deletion profiles observed.

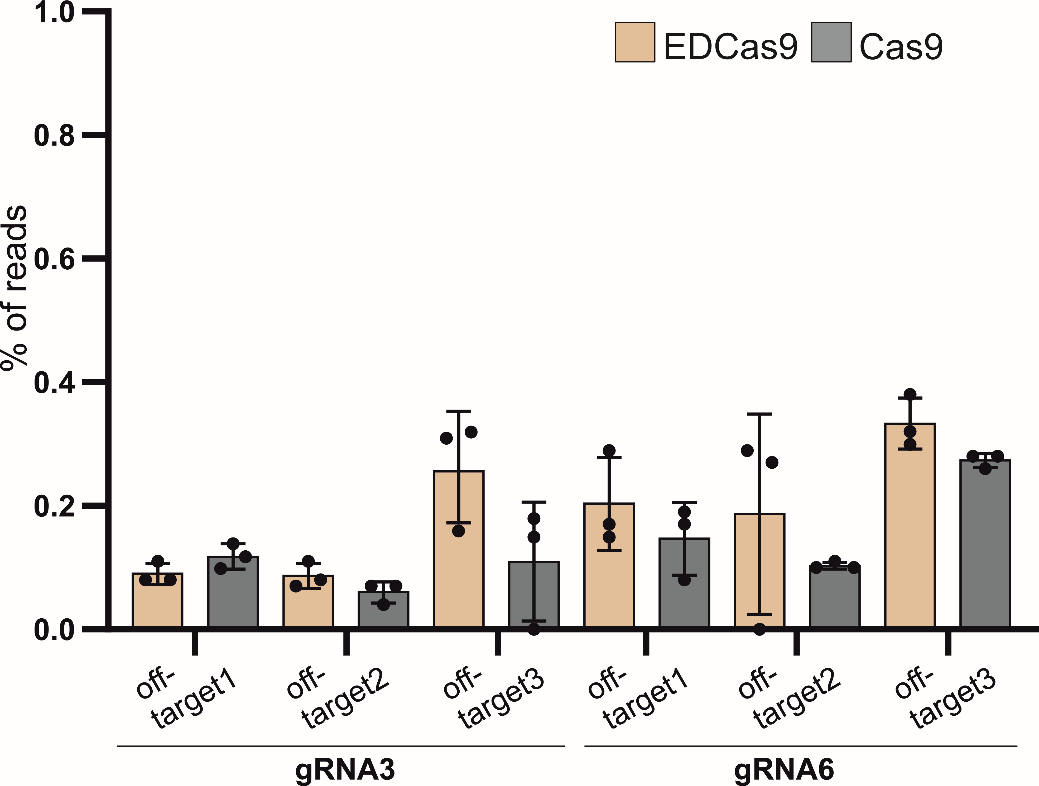

**Supplementary Figure 7: Quantification of the selected off-target sites for gRNA3 and gRNA6 coupled to EDCas9 and Cas9 in homozygous USH2A:c.7595-2144G patient-derived fibroblasts.** High-throughput sequencing was used to assess off-target potential. Results are depicted as percentage (%) of edited reads over the total number of reads ± standard deviation. Three (n=3) independent replicates were performed. No significant difference between EDCas9- and Cas9-treated cells is observed.

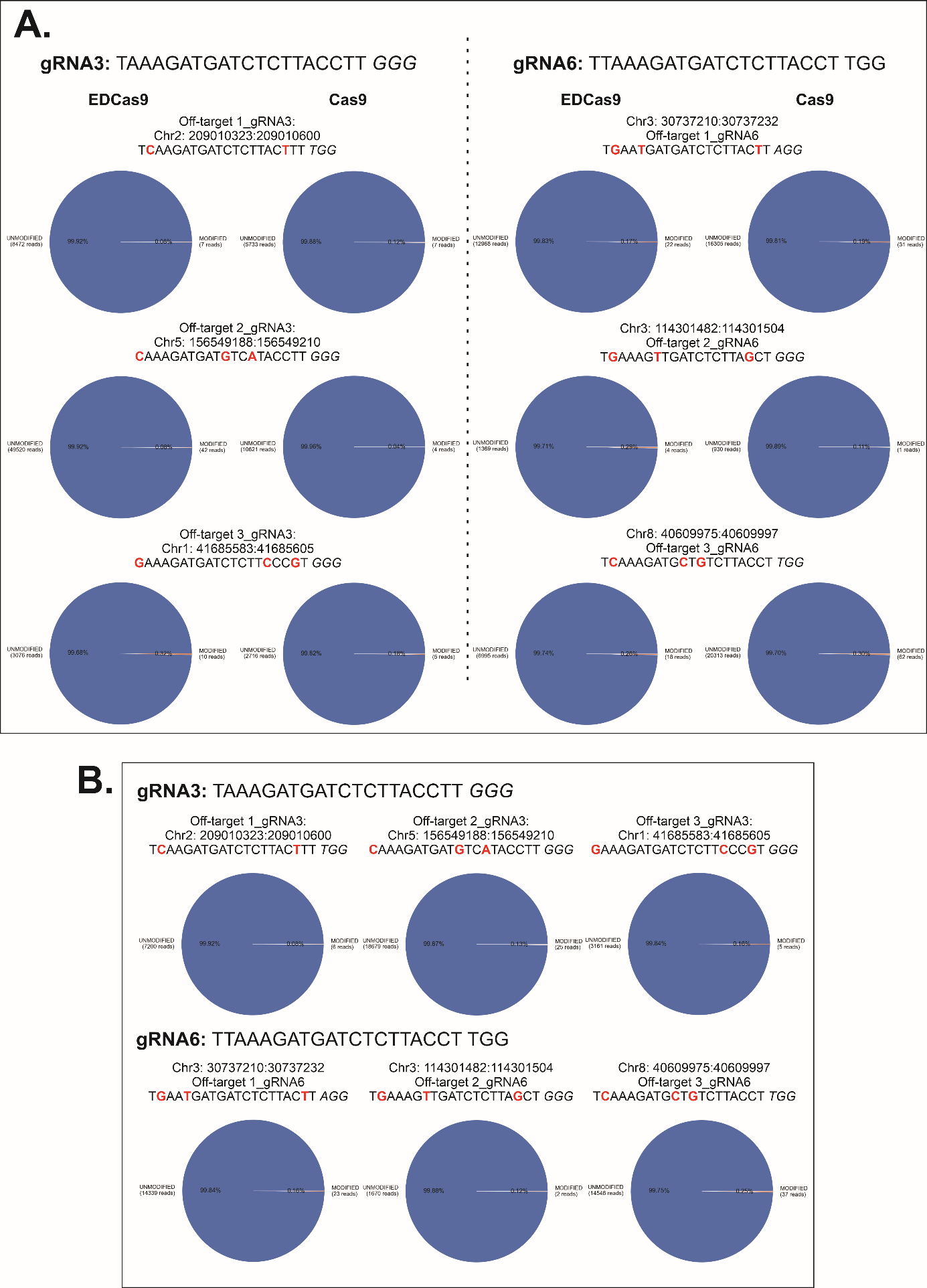

**Supplementary Figure 8: Representative examples of histograms depicting the amount of edited (blue) and non-edited (orange) reads for the selected off-target sites analyzed by high-throughput sequencing. (A)** Off-target analysis on patient-derived fibroblasts treated with gRNA3 and gRNA6 coupled to EDCas9 or Cas9. (**B**) Off-target analysis on non-treated patient-derived fibroblasts. (**A**,**B**) Location of the off-target sequence (chromosome: position) and sequence are reported above the histogram. In bold red the nucleotides mismatching with the gRNA sequence. In italics, the PAM sequence.
